# Blm Helicase Facilitates Rapid Replication of Repetitive DNA Sequences in early *Drosophila* Development

**DOI:** 10.1101/2021.06.16.448677

**Authors:** Jolee M. Ruchert, Morgan M Brady, Susan McMahan, Karly J. Lacey, Leigh C. Latta, Jeff Sekelsky, Eric P. Stoffregen

## Abstract

The absence of functional BLM DNA helicase, a member of the RecQ family of helicases, is responsible for the rare human disorder Bloom Syndrome, which results in developmental abnormalities, DNA repair defects, genomic instability, and a predisposition to cancer. In *Drosophila melanogaster*, the orthologous Blm protein is essential during early development when the embryo is under the control of maternal gene products. We show that lack of functional maternal Blm during the syncytial cell cycles of *Drosophila* embryonic development results in severe nuclear defects and lethality. Amongst the small fraction of embryos from *Blm* mutant mothers that survive to adulthood, a prominent sex-bias favors the class that inherits less repetitive DNA content, which serves as an endogenous source of replication stress. This selection against repetitive DNA content reflects a role for Blm in facilitating replication through repetitive sequences during the rapid S-phases of syncytial cell cycles. During these syncytial cycles, Blm is not required for complex DNA double-strand break repair; however, the progeny sex-bias resulting from the absence of maternal Blm is exacerbated by repetitive DNA sequences and by the slowing of replication fork progression, suggesting that the essential role for Blm during this stage is to manage replication fork stress brought about by impediments to fork progression. Additionally, our data suggest that Blm is only required to manage this replication stress during embryonic development, and likely only during the early, rapid syncytial cell cycles, and not at later developmental stages. These results provide novel insights into Blm function throughout development.

## INTRODUCTION

BLM DNA helicase, a member of the RecQ family of ATP-dependent helicases (Ellis, 1995), gives rise to Bloom Syndrome when absent in humans. Bloom Syndrome is characterized primarily by an increased risk of developing a broad range of cancers, but low birth weight, proportional dwarfism, photosensitive skin lesions, and premature aging are also commonly observed (German 1993; Chu and Hickson 2009; Kamenisch and Berneburg 2009). BLM plays multiple roles in the repair of DNA double strand breaks (DSBs) in both mitotic and meiotic cells (Adams *et al*. 2003; Bernstein *et al*. 2010; Croteau *et al*. 2014). Additionally, BLM is involved in the response to replication stress, defined here as a slowing or stalling of replication fork progression (Zeman and Cimprich 2013), which is caused by blocked, stalled, or collapsed replication forks (Davalos *et al*. 2004; Wu 2007). *Drosophila melanogaster* makes a compelling model for the study of BLM function due to the well-documented functional conservation between human BLM and *Drosophila melanogaster* Blm proteins with respect to their roles in DNA DSB repair and replication fork management (Karow *et al*. 2000; van Brabant *et al*. 2000; Machwe *et al*. 2005; Bachrati *et al*. 2006; McVey *et al*. 2007).

When initially identified in *Drosophila* as *mus309*, a mutagen-sensitive mutant allele on the third chromosome, *Blm* mutations were found to cause almost complete sterility in females (Boyd *et al*. 1981). Later analysis of this sterility defect determined the cause to be a maternal-effect embryonic lethality; the vast majority of embryos from *Blm* mothers do not hatch into larvae (McVey *et al*. 2007). Early embryonic development in *Drosophila* occurs in a syncytium; the syncytial embryonic cycles are the first 13 cell cycles occurring post-fertilization. The nuclei in the syncytium undergo rapid, mostly synchronous rounds of DNA synthesis (S) and mitosis (M) with no intervening gap phases (Foe and Alberts 1983). These syncytial cycles are under maternal control, with maternally deposited gene products (mRNA transcripts and protein) driving cellular processes until zygotic transcripts are produced in sufficient quantities. This transition between maternal and zygotic control occurs gradually from the late syncytial cycles until completion by the mid-blastula stage during cycle 14 (reviewed in Tadros and Lipshitz 2009; Kotadia *et al*. 2010; Laver *et al*. 2015). *Blm* mothers, therefore, fail to provision their eggs with maternal Blm products, resulting in embryos that lack functional Blm helicase. These embryos display an elevated incidence of anaphase bridge formation and other markers of DNA damage during syncytial cycles, as well as a subsequent low embryo hatch rate (McVey *et al*. 2007). These data led to the hypothesis that Blm is essential during the syncytial embryonic cell cycles.

Observations have been made of a sex-bias towards female progeny amongst the small percentage of total progeny from *Blm* mothers that are able to survive to adulthood (J. Sekelsky, K. P. Kohl, unpublished observations). One characteristic that differentiates early male and female embryos is the amount of repetitive DNA content in their respective genomes, with female genomes containing ~ 20 Mb less repetitive content than male genomes (Hoskins *et al*. 2002; Celniker and Rubin 2003; Brown *et al*. 2020). One possible explanation for the underrepresentation of male progeny from *Blm* mothers is that the primary role for Blm protein during early syncytial cell cycles is to manage replication challenges, such as those posed by replication fork stalling through repetitive DNA sequences. In this case, the extra repetitive sequence content in the male genome puts those embryos at a survival disadvantage.

Repetitive DNA sequences have been shown to cause replication fork pausing both *in vitro*(Kang *et al*. 1995; Gacy *et al*. 1998) and *in vivo* (Samadashwily *et al*. 1997; Ohshima *et al*. 1998). Repetitive DNA sequences can form several types of secondary structure that impede the replicative helicase (reviewed in Mirkin and Mirkin 2007), and members of the RecQ helicase family can unwind many of these types of secondary structure (reviewed in Sharma 2011). Replication fork pausing may be especially problematic in the extremely rapid syncytial S-phases. During the early syncytial cycles, the embryo replicates the entire genome in an estimated 3-4 minutes (Blumenthal *et al*. 1974), which lengthens in the late syncytial cycles (9-13) to between 5 and 14 minutes before lengthening considerably to 50 minutes at cycle 14 (Shermoen *et al*. 2010).

Taken together, these data suggest that Blm DNA helicase plays an essential role during syncytial embryonic development in *Drosophila*. More specifically, it suggests that the precise function of Blm may be in responding to replication stress during the rapid syncytial cell cycles. However, this proposed function of Blm has not been directly tested.

To begin our investigation, we collected examples of the severity of DNA damage that arises in embryos from *Blm* mothers. We then tested the hypothesis that Blm prevents such DNA damage during early embryonic development by responding to endogenous sources of replication stress, such as those posed by repetitive DNA sequences, by altering the amount of repetitive DNA content that was inherited by either male or female progeny from various crosses. This allowed us to determine whether the relative amount of repetitive DNA content in embryos that lack maternal Blm (those from *Blm* mothers) was correlated with survival to adulthood. If so, it would manifest as a sex-bias in the progeny favoring the class of flies that inherits less repetitive sequence content. We then tested whether a resulting progeny sex-bias is due to defects in the ability of Blm to repair DNA DSBs or whether the phenotype is correlated with a further slowing of replication fork progression. Lastly, we tested whether this essential role of Blm is restricted to syncytial cell cycles or whether this function(s) of Blm is also required in later developmental stages.

## RESULTS

### Embryos without maternal Blm exhibit significant DNA damage

Embryos that lack maternal Blm frequently form anaphase bridges during syncytial cycles (McVey *et al*. 2007). To further characterize the DNA damage phenotype of embryos from *Blm* mothers, we stained and fixed embryos with the DNA dye DAPI, an antibody to phosphorylated serine 10 of histone H3 (H3S10P, a marker of mitosis), and a phosphotyrosine (pY) antibody, which marks actin-rich cages surrounding syncytial nuclei. In embryos from wild-type mothers (*Blm*^+^), nuclei go through mitosis synchronously, and the actin-rich cages form a regular hexameric array (Fig 1A). In an embryo in which most nuclei are in mitosis, there are sometimes examples of nuclei that have not entered mitosis (Fig. 1A, arrows) or of cages that appear to lack a nucleus (Fig. 1A, dotted outline). This latter phenotype stems from nuclear damage during the rapid S and M cycles. At the syncytial blastoderm stage, nuclei that have sustained sufficient damage exit the cell cycle and drop out of the cortex into the interior of the embryo, where they will not contribute to the embryo proper (Sullivan *et al*. 1993; O’Dor *et al*. 2006). This process, termed nuclear fallout, sometimes involves several adjacent nuclei, likely the descendants of a single nucleus that suffered damage in an earlier cell cycle.

**Figure 1.**
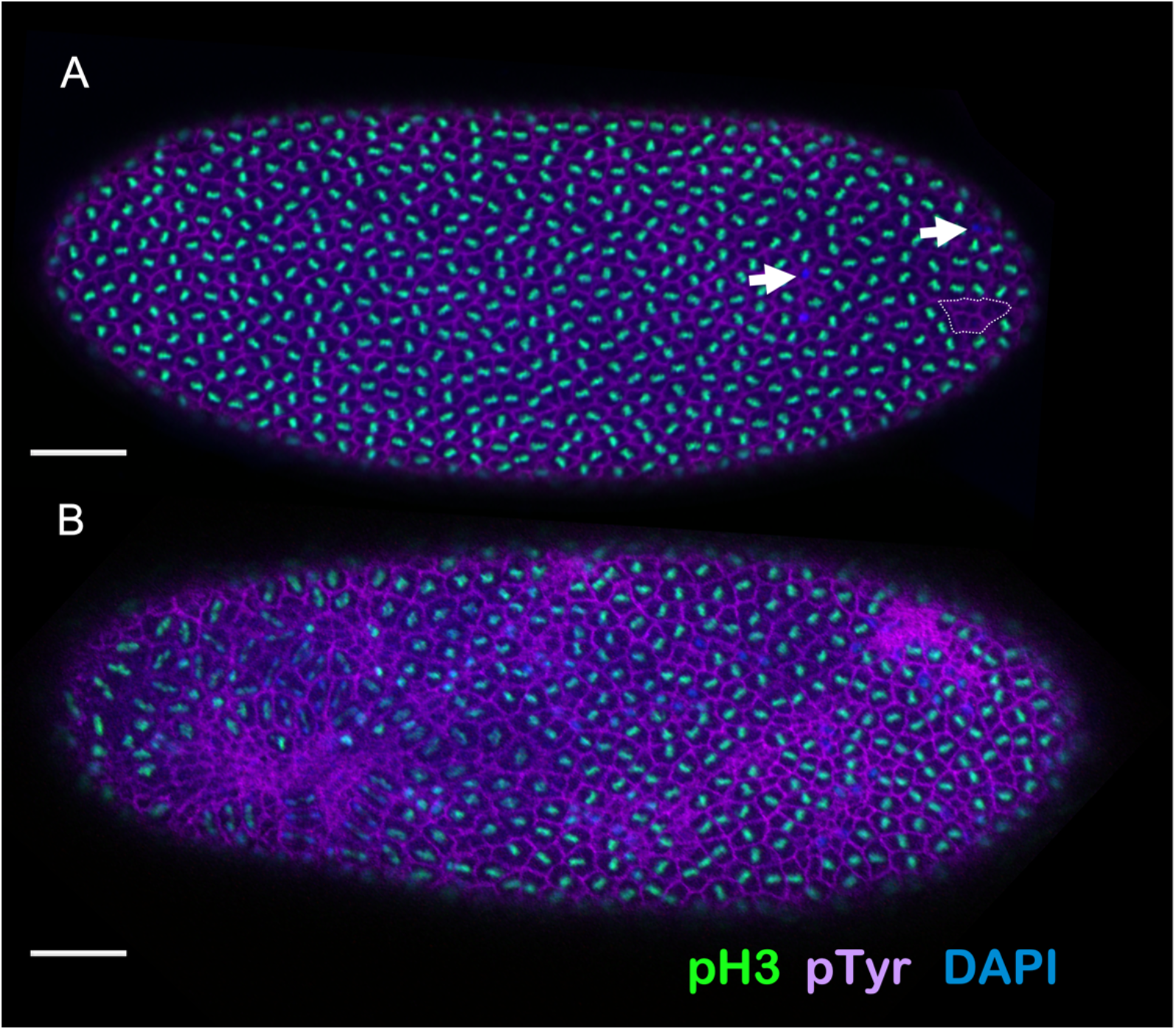
Early embryos lacking maternal Blm display severe nuclear defects. **(A)** An example of a late (nuclear cycle 11-13) syncytial stage embryo from a mother wild-type for *Blm* (*w^1118^*), fixed and stained with the DNA dye DAPI (blue) and antibodies to phosphotyrosine (purple; labels actin cages surrounding nuclei) and to phosphorylated serine-10 of histone H3 (pH3; green). Nearly uniform labeling of pH3 shows that mitosis is synchronous across the embryo. A few instances of nuclei with DNA but no pH3 (arrows) indicate instances of asynchrony or other nuclear defects. One cluster of empty actin cages (dotted outline) reveals a region of nuclear fallout. **(B)** A late (nuclear cycle 11-13) syncytial stage embryo from a *Blm* mother, fixed and stained as in (A). Massive asynchrony and nuclear fallout are evident. Bar, 50 μm.

Severe defects are observed amongst embryos that develop without maternal Blm helicase (embryos from *Blm* mothers; *Blm*^-^) by cycle 14; these defective embryos exhibit large patches of asynchrony, nuclear fallout, and large swaths of collapsed actin cages (Fig. 1B). This extreme level of nuclear damage likely accounts for the non-viability of most embryos from *Blm* mothers, suggesting that maternally deposited Blm has one or more essential functions during the rapid syncytial cell cycles that prevent DNA damage-induced nuclear fallout and the subsequent failure-to-hatch of Blm-null embryos.

### Survival of progeny from *Blm* mothers is inversely correlated with repetitive DNA sequence load

Although most embryos that lack maternal Blm die prior to the larval stages of development (McVey *et al*. 2007), a small percentage survive to adulthood. Among the adult survivors, there is an observed bias toward female progeny compared to male progeny that we set out to quantify. First, we tested a condition in which both female and male flies had normal sex chromosomes (*XX* for mothers and *XY* for fathers) and progeny inherited normal sex chromosomes (*XX* for female progeny and *XY* for male progeny) (Fig. 2A). Under these conditions, when females lacking functional Blm (*Blm* females) were crossed to males with wild-type *Blm* (we used *w^1118^* as our *Blm*^+^ females and for our males with normal sex chromosomes), more than 70% of the adult progeny were female, which is significantly different from the expected 50% female ratio (Fig. 2A, t = 13.041, df = 50, p-value < 0.0001) and significantly different than we saw when *w^1118^* male flies were crossed to *Blm*^+^ females (50.1% female progeny; t = 11.49, df = 71.706, *p* < 0.0001). One possible explanation for this female sex-bias amongst the progeny from *Blm* mothers is that mutations accumulate in germline cells of the *Blm* mutant mother, resulting in non-viability of sons that receive a maternal *X* chromosome with one or more lethal mutations. However, since syncytial development runs almost entirely on maternal gene products with very little contribution from the zygotic genotype (reviewed in Tadros and Lipshitz 2009; Kotadia *et al*. 2010; Laver *et al*. 2015), this scenario seemed unlikely to account for early embryonic defects. Therefore, we considered the possibility that it is the presence of the *Y* chromosome that makes male embryos less likely to survive. We used a series of *X* and *Y* chromosomal rearrangements to test this hypothesis.

**Figure 2.**
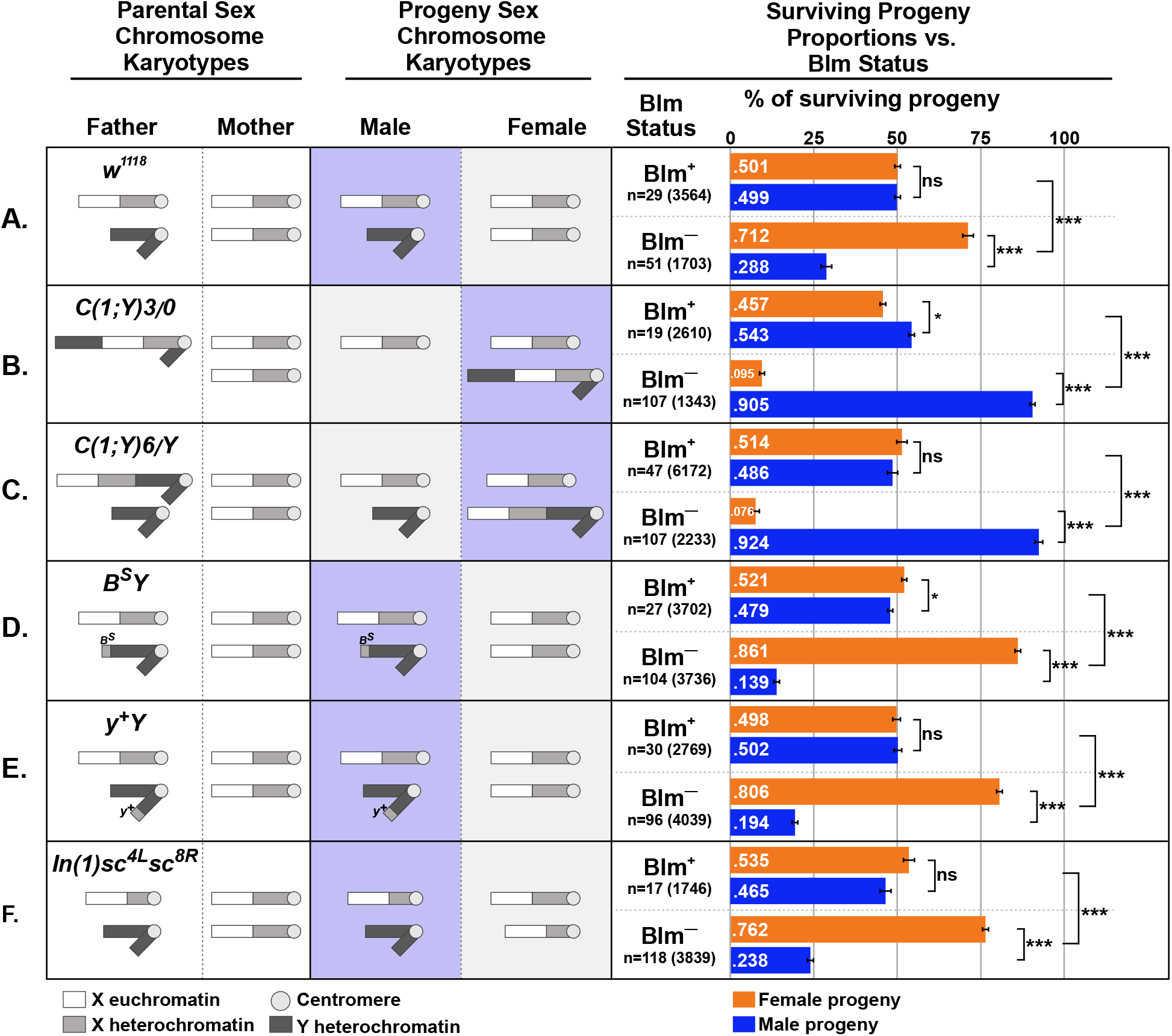
Survival of progeny from *Blm* mothers is inversely correlated with repetitive sequence load. For each cross (A-F), the sex chromosome karyotypes of the father and mother, with the approximate areas of euchromatin or heterochromatin content on the *X* or *Y* chromosomes indicated by white (euchromatin), light gray (*X* chromosome heterochromatin), or dark gray (*Y* chromosome heterochromatin) are shown. For all crosses, the mothers, whether *Blm* (fail to provide functional Blm protein to their eggs, depicted as *Blm*^−^) or *w^1118^* (provide functional Blm protein to their eggs, depicted as *Blm*^+^), have normal sex chromosomes (two normal *X* chromosomes). See text for more details on the chromosomal content for each male. Next are depictions of the sex chromosome karyotypes of the progeny from each cross (not accounting for progeny classes resulting from chromosomal nondisjunction). The progeny class from each cross that inherits more repetitive DNA sequence content is indicated by the purple-boxed chromosomes. Lastly, for each cross, the relative proportions are shown of surviving male (blue bars) and female (orange bars) flies for each cross to mothers with either functional (Blm^+^) or nonfunctional (Blm^-^) Blm packaged into the eggs. The proportion of female versus male progeny was recorded for each individual experiment (vial), and those proportions were averaged. The total number of experiments (n) as well as the total number of flies scored for each condition is shown. In all crosses, there is a progeny sex-bias favoring the progeny class that inherits less repetitive DNA content. Error bars represent 2 × SEM. ns – not significantly different (p > .05); * p < 0.05; *** p < .0001. All p-values were calculated with t-tests comparing the observed proportion to an expected proportion of 0.50 or by comparing two observed proportions of female progeny to one another.

We crossed *Blm* females to males carrying a compound chromosome that fuses the *X* and *Y*, designated *C(1;Y)*. We first used *C(1;Y)3*/O males (“O” indicates the lack of a second sex chromosome). These males produce sperm that carry either the *C(1;Y)3* chromosome or no sex chromosome; when sperm from these males fertilize a normal egg containing a single *X* chromosome, the result is either a *C(1;Y)3/X* zygote or an *X*/O zygote. In *Drosophila*, sex is determined by the ratio of X:autosomes and not by the presence of a *Y* chromosome (Salz and Erickson 2010); therefore, the *X/XY* sex chromosome karyotype of the *C(1;Y)3/X* zygote will develop as female and the X/O zygote will develop as male. In this cross, more than 90% of the surviving progeny were male (*X*/O), which differs significantly from the results when *C(1;Y)3*/O males were crossed to *Blm*^+^ females (Fig. 2B, t = 18.418, df = 40.202, p < 0.0001). We did a second cross with an independently generated compound chromosome, *C(1;Y)6*. In this cross, fathers were *C(1;Y)6*/Y, and resulting progeny were either *X/XY* females or *X/Y* males. Again, more than 90% of the progeny surviving to adulthood were male (this time with *X/Y* genotype), which was significantly different from what is seen when the *C(1;Y)6*/*Y* males are crossed to *Blm*^+^ females (Fig. 2C, t =40.441, df = 93.376, p < 0.0001). This data suggests that the sex-bias seen in the progeny from *Blm* mothers is not caused by problems with the maternally derived *X* chromosome; however, the sex-bias does correlate with the presence of a *Y* chromosome (although not to the male sex specifically) and/or with increased repetitive DNA content when both classes of progeny inherit a *Y* chromosome (in the case of *X/Y* vs. *X/XY* progeny the difference is not the presence of the *Y* but the additional X chromosome).

The *Drosophila melanogaster X* and *Y* chromosomes are each about 40 Mb, but the *Y* is largely composed of highly repetitive satellite DNA sequences, whereas less than 50% of the *X* is highly repetitive (Hoskins *et al*. 2002; Celniker and Rubin 2003; Brown *et al*. 2020). Thus, although *X/X* females and *X/Y* males have similar amounts of DNA per cell, *X/Y* individuals have far more sex chromosome satellite DNA (~60 Mb) than *X/X* siblings (~40 Mb). The precise DNA content of *C(1;Y)6* and *C(1;Y)3* are unknown, but, in crosses involving either male, there would be a greater amount of satellite sequences in *C(1)Y/X* female progeny than in *X/O* or *X/Y* male progeny. Direct comparisons between the crosses involving either *C(1;Y)3* or *C(1;Y)6* males are difficult to make due to the differences in both the inherited *C(1;Y)* chromosome in females and the difference in the presence or absence of a *Y* chromosome in the male progeny. We would expect that if the inherited *C(1;Y)* chromosome were the same in the female progeny from both *C(1;Y)3* and *C(1;Y)6* males, then the male progeny sex-bias would be more pronounced in the *C(1;Y)3* cross due to the increased probability of survival of the *X*/O males from that cross compared with the *X/Y* males from the cross with *C(1;Y)6* males. However, the lack of a substantial difference between the male progeny sex-bias between these crosses could be due to differences in the *C(1;Y)* chromosomes and/or to the relative difficulty in recovering progeny from the cross with *C(1;Y)3* males (which could also be caused by this particular *C(1;Y)* chromosome).

The results described above suggested that poor survival of embryos lacking maternal Blm helicase is related to increased dosage of one or more satellite sequences. To test this hypothesis, we crossed *Blm* females to males with either of two *Y* chromosome derivatives, *Dp(1;Y)B^S^* (hereafter called *B^S^Y*) and *Dp(1;Y)y*^+^ (*y*^+^*Y*). These chromosomes each contain the normal *Y* chromosome DNA sequence contribution plus an additional amount of *X*-derived satellite sequence (Gatti and Pimpinelli 1983). In both cases, the surviving progeny were significantly biased towards females compared to the expected 50% (t = 29.49, df = 93.556, p < 0.0001 for *B^S^Y*, Fig. 2D; t = 21.624, df = 63.984 p < 0.0001 for *y*^+^*Y*, Fig. 2E) and significantly more female-biased than that seen in crosses involving males with a normal *Y* chromosome (86.1% and 80.1% female progeny from *B^S^Y* and *y*^+^*Y* fathers, respectively, compared to 71.2% female progeny from *w^1118^* fathers, t = 8.0738, df = 78.557 and t = 5.1182, df = 77.964 respectively, p < 0.0001 for both comparisons). These results are consistent with the hypothesis that additional repetitive sequence content correlates with reduced embryonic survival.

We also reduced the amount of *X* satellite sequence packaged into female embryos by crossing *Blm* females to males with a normal *Y* chromosome and an *X* chromosome carrying *In(1)sc^4L^sc^8R^*, a rearrangement that deletes about 2/3 of the *X* satellite sequences. Daughters from this cross have less sex chromosome repetitive DNA sequences than do their male siblings, and less than female progeny from crosses involving males with a normal *X* chromosome. In support of our hypothesis, the female sex-bias in surviving progeny was more pronounced in the cross with *In(1)sc^4L^sc^8R^* males (Fig. 2F) than with normal *XY* males (Fig. 2A) (t = 2.6689, df = 81.915, p = 0.009172). Taken together, our data indicate an inverse relationship between sex chromosome repetitive DNA sequence content and survival to adulthood in embryos lacking maternal Blm.

### The progeny sex-bias is established during embryonic development

Embryos from *Blm* mothers exhibit severe nuclear defects during syncytial embryonic development, and progeny that survive the Blm-null embryonic environment exhibit a female sex-bias when progeny inherit normal sex chromosomes (*XX* for females and *XY* for males). However, it was unknown whether this sex-bias occurs due to differential survival of *XX* embryos or whether the survival differential occurs later in development (during the larval stages, for example). To address this question, we analyzed embryos, larvae, and adults to see when in development the sex-bias first manifests. To study embryos, *Blm* mothers that express eGFP under an *Sxl* promoter that is only transcribed in embryos with a 1:1 ratio of *X* chromosomes to autosomes (Schutt and Nothiger 2000; Thompson *et al*. 2004), were crossed to *w^1118^* males. The result of this is eGFP expression in *XX* or *XXY* embryos but not in *XY* or *XO* embryos. In embryos imaged 6-8 hours into development, no significant difference was detected in the numbers of female versus male embryos (Fig. 3).

**Figure 3:**
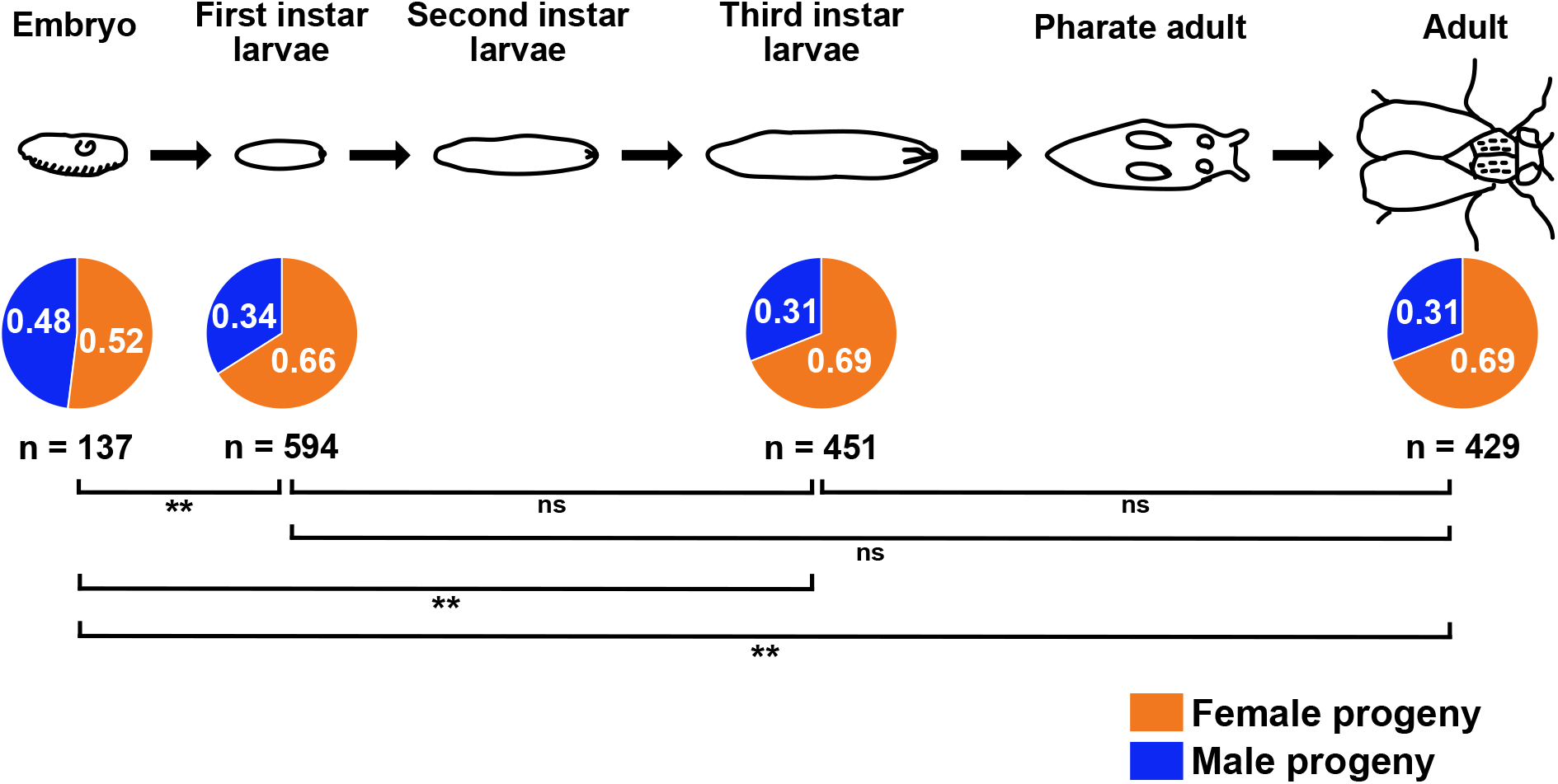
The sex-bias in progeny from *Blm* mothers is established during embryonic development. Embryos sexed 6-8 hours into development (past the syncytial stages, but as early as we could detect the eGFP expressed via the *Sxl* promoter and determine sex) showed no evidence of a sex-bias; however, in a separate set of experiments, 1^st^ instar larvae showed a female sex-bias that did not significantly change throughout the remainder of development (3^rd^ instar larvae and adults from the same embryo collections that provided the 1 ^st^ instar larvae data showed no difference in the proportion that were female). The female sex-bias that is evident at the 1^st^ and 3^rd^ instar larval stages, and in surviving adults, is not significantly different from one stage of development to another, but all are significantly different from the proportion of embryos that are female (the proportion of female embryos is not significantly different from the expected 50:50 ratio). ns = not significant (p > .05); ** p < .01; ** p < .001. All p-values were calculated using two-tailed z-tests to compare the observed proportions to one another.

We next assessed the ratio of female versus male larvae at the first and third instar larval stages, as well as the ratio of female versus male adults from the same pool of collected embryos. For this assay, we marked *X* chromosomes with a *y^1^* allele; this mutation of the *yellow* gene results in brown larval mouth hooks instead of black without a wild-type copy of *y* (*y*^+^). When *y^1^*; *Blm* females were crossed to *w^1118^* males (with a *y^+^* on their *X* chromosome), all surviving female larvae are *y*^+^ / *y^1^*, and thus have the wild-type black mouth hooks. Meanwhile, male larvae are hemizygous for the *y^1^* allele and have brown mouth hooks. At the first instar larval stage there is a clear female sex-bias, which does not change throughout the remainder of development (Fig. 3). The proportion of embryos that were female in the preceding experiment (0.52) is significantly different from the proportion of female larvae at the first and third instar stages (0.66 and 0.69, respectively; z = −3.1416, p = 0.00168 and z = −3.7332 p = 0.0002, respectively) and from the proportion of female adults (0.69, z = −3.7191, p < 0.001). There is no significant difference between the proportions of female progeny when comparing the larval stages to one another or to the adult stage (1^st^ instar vs. 3^rd^ instar, z = −1.0314, p = 0.30; 1^st^ instar vs. adult, z = −1.0338, p = 0.30; 3^rd^ instar vs. adult, z = −0.0164, p = 0.98), indicating that the female sex-bias (or the bias favoring the class of progeny with less repetitive DNA sequence content) is established by the time of egg hatching.

### The essential role for Blm is restricted to early embryogenesis

*Blm* mothers display a severe maternal-effect lethality (McVey *et al*. 2007); however, when not exposed to DNA damage, *Blm* flies survive throughout the stages of development that rely on zygotic-, larval-, and adult-produced Blm protein. This led us to hypothesize that the essential role for Blm is restricted to the early syncytial cell cycles, which are dependent upon maternally derived Blm products. To test this hypothesis, we crossed *Blm* females to males that were heterozygous for *Blm*. Half of the resulting progeny have one functional copy of *Blm* and can produce functional Blm protein upon activation of zygotic transcription, while the other half never produce functional Blm protein. No difference in survival to adulthood was detected between progeny that were either homozygous or compound heterozygous for *Blm* mutant alleles (unable to produce functional Blm after initiation of zygotic control) versus those that were heterozygous for *Blm* (produce functional Blm from one allele after initiation of zygotic control) (Fig. 4). This data indicates that around the time zygotic transcription of *Blm* begins, the protein is no longer a significant determining factor in survival to adulthood.

**Figure 4:**
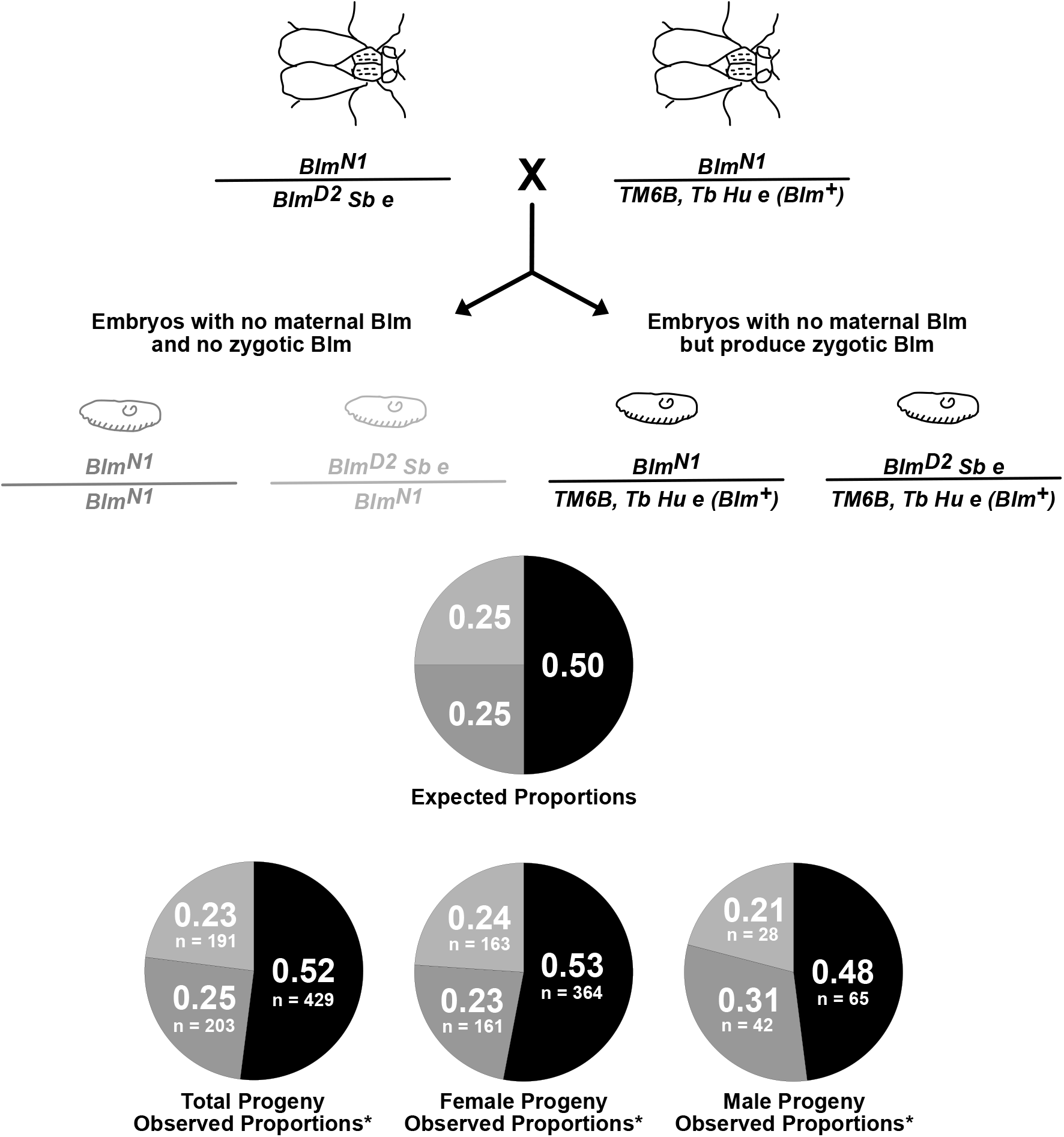
The essential role for Blm is restricted to embryogenesis. *Blm* mutant females (*Blm^N1^* / *Blm^D2^ Sb e*) were crossed to males that were heterozygous for *Blm (Blm^N1^* / *TM6B, Tb Hu e*). All resulting embryos develop without maternal Blm protein. However, half of the progeny will have functional Blm beginning with the onset of zygotic transcription (right, embryo and genotype shown in black) and half of the progeny will never produce functional Blm (left, light and dark gray embryos and genotypes). Surviving adult progeny display phenotypic markers that allowed them to be classified as either heterozygous (a *Blm*^+^ allele along with either the *Blm^N1^* or *Blm^D2^* null allele), compound heterozygous (both the *Blm^N1^* and *Blm^D2^* null alleles), or homozygous (two *Blm^N1^* alleles). We wanted to differentiate between the *Blm*-null progeny *(Blm^N1^* / *BlmD^2^ Sb e* versus *Blm^N1^* / *Blm^N1^*) to ensure that survival was not impacted by potential off-site mutations in the homozygous class. The top pie chart shows the expected proportions of surviving progeny for each genotypic class, with the pie slice color corresponding to the same-colored genotype shown above. The bottom row of pie charts shows the observed proportions of each genotypic class in total progeny and in female and male progeny, with the number of progeny counted for each class indicated. All values are consistent with expected Mendelian ratios (for total progeny, χ^2^ = 0.9745, p = .614; for female progeny, χ2 = 1.1701, p = .557; for male progeny, χ^2^ = 1.4531, p = .484).

### Heterochromatin localization is not temporally altered in embryos from *Blm* mothers

The data quantifying the progeny sex-biases observed when males containing variable repetitive DNA sequence content on their sex chromosomes are crossed to *Blm* mothers (Fig. 2) supports our hypothesis that Blm is necessary during syncytial cycles to respond to the challenges of replicating through repetitive DNA sequences. However, we wanted to rule out other potential causes of the embryonic DNA damage and subsequent progeny sex-bias. First, we investigated whether early changes to heterochromatin structure might play a role. Markers of heterochromatin structure, such as HP1 and corresponding repressive histone modifications, are usually not observed in wild-type embryos until cycle 14 of the syncytial embryo (Shermoen *et al*. 2010), which is later in development than DNA damage in embryos from *Blm* mutant mothers is detected (McVey *et al*. 2007 and Fig. 1). To test whether the sex-bias in progeny from *Blm* mothers is related to a change in the temporal pattern of heterochromatin formation, we utilized an antibody against H3K9me2 and confirmed that embryos from both *Blm* and *Blm*^+^ (*w^1118^*) mothers lack H3K9me2 localization until after cellularization occurs at cycle 14 (Figure 5). Since embryos from *Blm* mothers do not have an altered temporal pattern of heterochromatin formation, it is unlikely that changes to heterochromatin structure are the cause of the nuclear defects observed.

**Figure 5:**
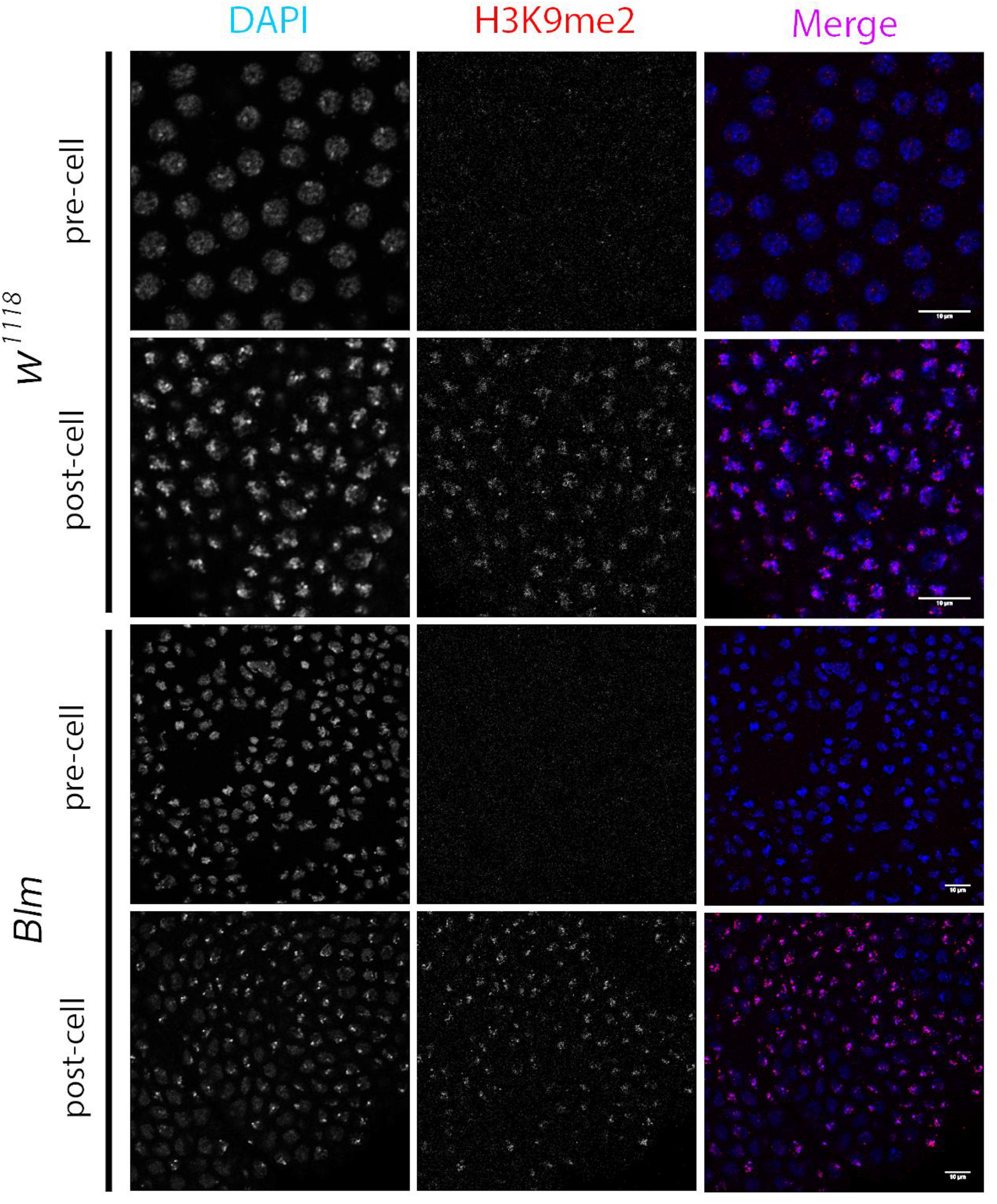
H3K9me2 localizes to nuclei following cellularization at nuclear cycle 14 in both wild-type and *Blm* mutant embryos. Wild-type (*w^1118^*) (top) and *Blm* (bottom) embryos were divided into pre-cellularization (pre-cell) or post-cellularization (post-cell) at nuclear cycle 14 (the end of syncytial development). The left column shows DNA (DAPI); middle column shows H3K9me2 (α-H3K9me2); right column shows a merged image (DNA, blue; H3K9me2, red). Scale bars represent 10 μm.

### The N-terminus of Blm, which participates in recombination and repair functions, is not essential for embryonic development

The next question we addressed was whether the nuclear damage and subsequent progeny sex-bias is related to an inability to properly repair DNA DSBs in the embryos from *Blm* mothers. The *Blm^N2^* mutant allele, an N-terminal deletion of the first 566 amino acids, is a partial separation-of-function mutant allele that is deficient in repair of DSBs; the *Blm^N2^* allele is null for SDSA and shows mitotic and meiotic recombination defects, but this allele retains its helicase domain (McVey *et al*. 2007). *Blm^N2^* mutants retain an ability to process stalled or blocked replication forks, as evidenced by the reduced maternal-effect lethality compared to *Blm*-null alleles (McVey *et al*. 2007) and the viability or increased survival of double mutants involving *Blm^N2^* and structure-selective endonucleases, such as *mus81*; *Blm^N2^* and *Gen Blm^N2^* mutants compared to double mutants involving *Blm*-null alleles (Trowbridge *et al*. 2007; Andersen *et al*. 2011). These endonucleases can provide an alternative resolution to double-Holliday junctions that form following DNA DSBs in the absence of Blm. These data suggest that the severe early lethality of embryos from *Blm* mothers may not be due to defects in complex DSB repair, but rather to a failure to properly respond to replication stress. To further test this hypothesis, we utilized the *Blm^N2^* allele to assess the progeny sex-bias correlation to variable repetitive DNA content, but under conditions where the DSB repair functions of Blm are impaired but the helicase domain remains intact.

As predicted, the progeny sex-bias is ameliorated in the surviving progeny from *Blm^N2^* mothers, as compared to mothers with two *Blm*-null alleles, when crossed to *w^1118^*, *B^S^Y*, or *C(1;Y)6* fathers (Fig. 6). In the crosses involving *w^1118^* fathers, the proportion of female progeny from *Blm^N2^* mothers was slightly elevated compared to the crosses to *Blm*^+^ mothers (t = 2.687, df = 122, p = 0.02). In the crosses to *B^S^Y* fathers, there is no significant difference in the proportion of female progeny from *Blm^N2^* and *Blm*^+^ mothers (t = 0.302, df = 166, p = 0.95). A less severe male progeny sex-bias was detected in the crosses between *Blm^N2^* females and *C(1;Y)6* males (significantly different from crosses to *Blm*^+^ mothers [t = 36.758, df = 190, p < 0.0001] but also significantly different from crosses to *Blm (Blm^N1/D2^, null*) mothers, [t = 24.778, df =190, p < 0.0001]). Taken together, these data support the hypothesis that the essential syncytial function of Blm is not related to complex DSB repair.

**Figure 6:**
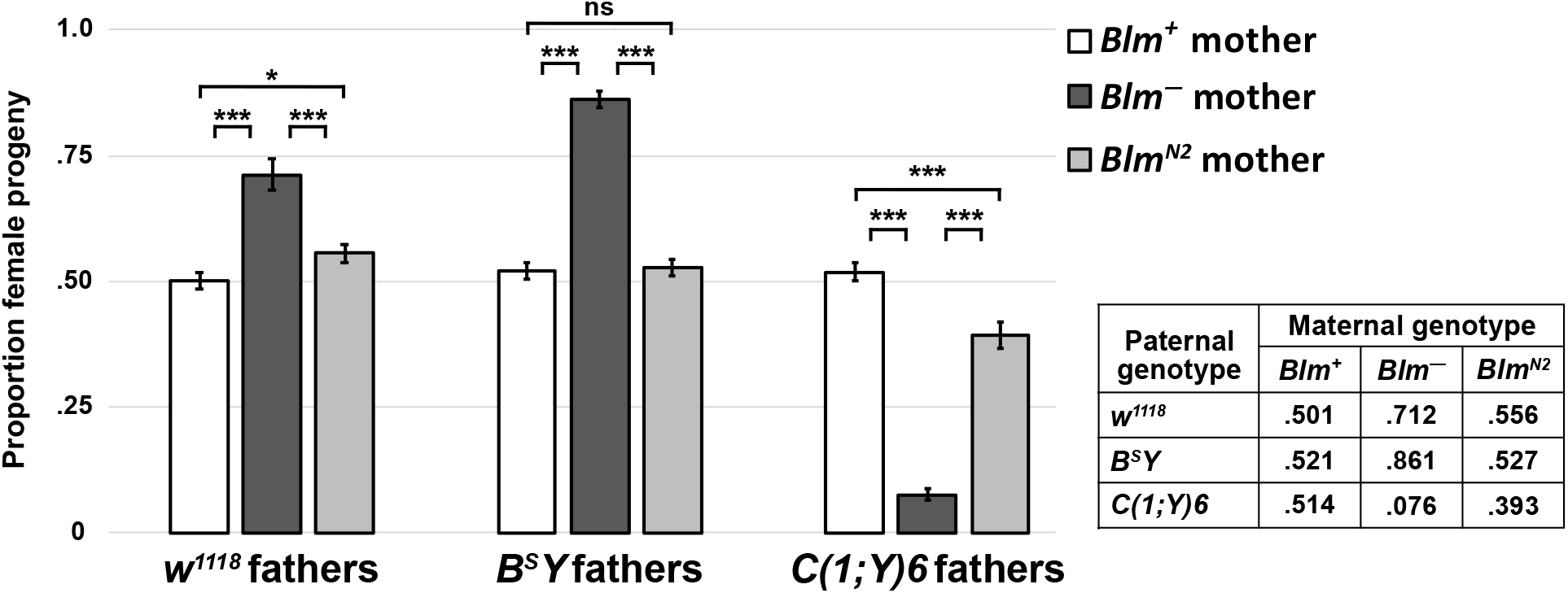
The sex-bias amongst progeny from *Blm* mothers is not due to DSB repair defects. The proportion of female progeny from *w^1118^* (*Blm*^+^) and *Blm* (*Blm*^-^) mothers (white and dark gray bars, respectively) is taken from Fig. 2 and compared to the proportion of female progeny from *Blm^N2^* mothers (light gray bars). *Blm*^-^ mothers carry two genetically null alleles of *Blm* (*Blm^N1^* and *Blm^D2^*). The *Blm^N2^* allele, however, codes for a protein product that is deficient for DSB repair but retains the entire helicase domain. In progeny from *Blm^N2^* mothers, who provide only this DSB repair-deficient protein to their embryos, the sex-bias favoring the progeny class that inherits less repetitive DNA is ameliorated. In all crosses, the proportion of female progeny from *Blm*^+^ and *Blm^N2^* mothers is significantly different than from *Blm*^-^ mothers (p < .0001 for all comparisons). In the crosses involving *B^S^Y* fathers, the proportion of female progeny from *Blm^N2^* mothers is not significantly different from the proportion from *Blm*^+^ mothers (p > .05). In the crosses involving *w^1118^* fathers, the proportion of female progeny from *Blm^N2^* mothers is only slightly different from that seen from *Blm*^+^ mothers (p < .05). In crosses involving *C(1:Y)6* fathers, the proportion of female progeny from *Blm^N2^* mothers is still significantly different from the proportion from *Blm*^+^ mothers (p < .0001), but this bias is far less pronounced than the bias amongst the progeny from *Blm*^-^ mothers. The number of replicates and total number of flies counted for *Blm^N2^* progeny were as follows: *w^1118^* fathers (n = 45; 5121); *B^S^Y* fathers (n = 38; 3577); *C(1;Y)6* fathers (n = 39; 3313). ns – not significantly different (p > .05); * p < .05; *** p < .0001. All p-values were calculated with one-way ANOVA tests followed by post-hoc comparisons of the observed proportions of female progeny to one another.

### Reducing Polα dosage exacerbates defects due to the absence of Blm

Since our data support a model by which the essential role for Blm in syncytial cycles is not to repair DSBs but rather to ensure proper replication through repetitive DNA sequences by responding to the replication stress induced by slowed or stalled replication forks, we hypothesized that slowing replication fork progression would further exacerbate the consequences posed by a lack of Blm protein during syncytial cycles. To test this hypothesis, we crossed males with variable repetitive DNA content on their sex chromosomes to either *w^1118^* (*Blm*^+^) or *Blm* mothers that contained only one functional copy of *DNA polymerase-α 180* (*Polα*), which codes for the catalytic subunit of the replicative polymerase. Genetically reducing Polα in this manner has been implicated in a slowing of DNA replication (LaRoque, 2007). This reduction of maternal Polα had no significant effect on the proportion of female progeny from *Blm*^+^ mothers (*w^1118^*; *Polα*^+/-^), who provide functional Blm protein to their eggs (Fig. 7, t = 0.512, df = 147, p =0.9561). However, in the absence of maternal Blm protein during embryogenesis, a genetic reduction of maternal Polα (eggs derived from *Blm Pola*^+/-^ mothers) results in a slight, but not statistically significant, exacerbation of the severe sex-bias seen in progeny from *Blm* mothers; with the sex-bias favoring the class of progeny that inherits less repetitive DNA, females in this case (Fig. 7, t = 1.979, df = 147, p = 0.2006). The male progeny sex-bias seen in the cross of *Blm Polα*^+/-^ mothers to *C(1;Y)6* males is significantly more pronounced than that from the cross to *Blm* mothers (t = 6.193, df = 212, p < 0.0001). In total, these data support the hypothesis that Blm is necessary to respond to replication stress caused by slowing or stalled replication during syncytial cycles.

**Figure 7:**
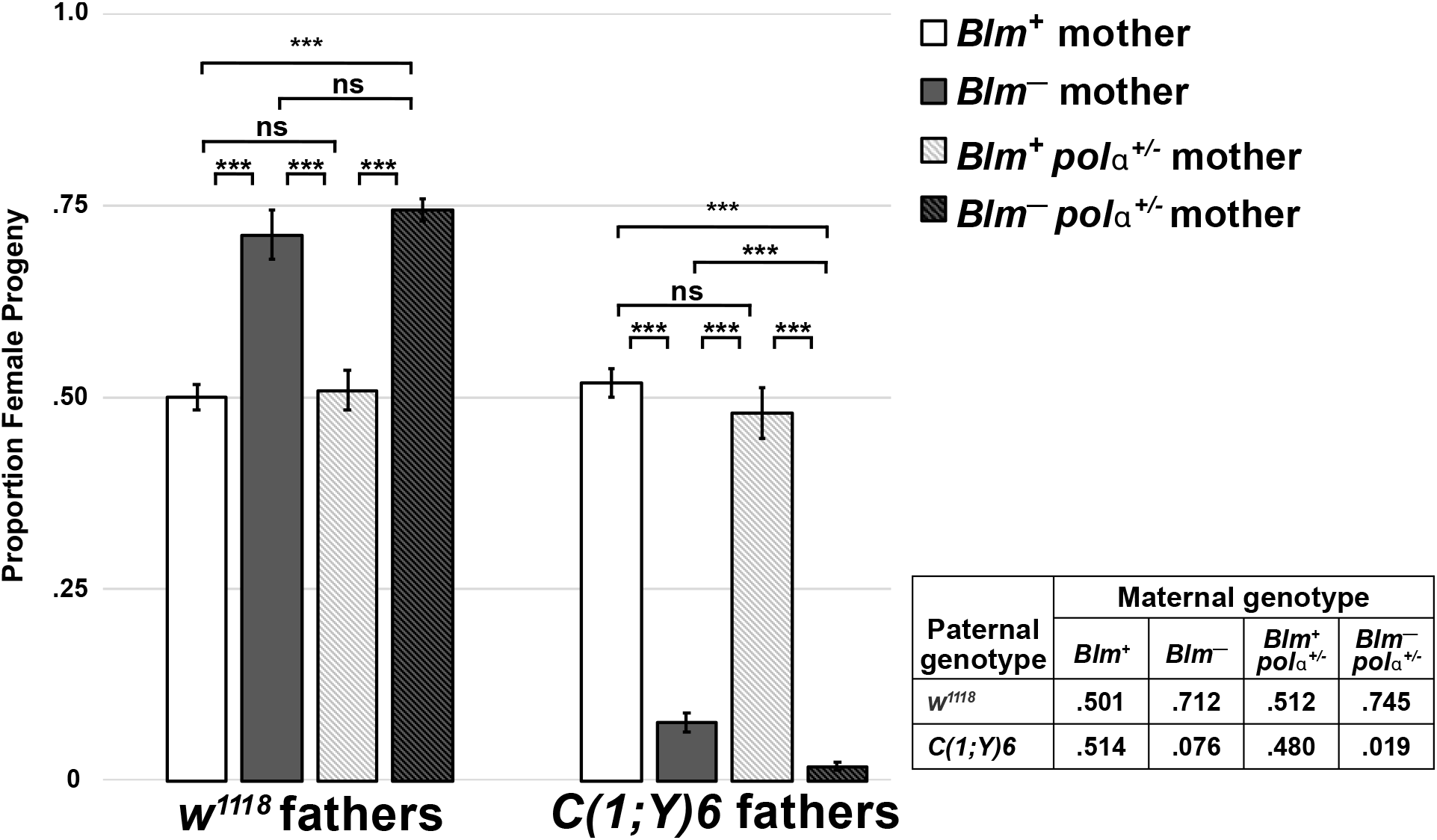
A genetic reduction of Polα exacerbates the sex-bias amongst progeny from *Blm*^-^ mothers. The proportion of female progeny from *w^1118^* (*Blm*^+^) and *Blm* (*Blm*^-^) mothers is taken from Fig. 2 and compared to *Blm*^+^ and *Blm*^-^ mothers that also package reduced Polα in their embryos (*Blm*^+^ *polα*^+/-^ and *Blm*^−^ *polα*^+/−^, respectively). In a *Blm*^+^ background, mothers who provide functional Blm to their eggs are not affected by a genetic reduction in Polα (no significant difference between *Blm*^+^ and *Blm*^+^ *polα*^+/−^ mothers in the progeny sex-bias in crosses to *w^1118^* or *C(1;Y)6* fathers; p > .05 for both comparisons). However, in a *Blm* background (*Blm*^−^ mothers), reducing Polα further exacerbates the significant progeny sex-bias that favors the class of flies that inherits less repetitive DNA content. In the crosses involving *w^1118^* fathers, the female progeny sex-bias was more pronounced, but did not reach statistical significance (p > .05). However, in crosses involving *C(1;Y)6* fathers, the exacerbation of the sex-bias favoring male progeny is significantly different compared to the progeny from *Blm*^-^ mothers (p < .0001). The number of replicates and total number of flies counted for *Blm^+^polα*^+/−^ progeny were as follows: *w^1118^* fathers (n = 25; 2439); *C(l;Y)6* fathers (n = 17; 1818). The number of replicates and total number of flies counted for *Blm polα’* progeny were as follows: *w^1118^* fathers (n = 46; 3265); *C(1;Y)6* fathers (n = 45; 2929). ns – not significantly different (p > .05); *** p < .0001. All p-values were calculated with one-way ANOVA tests followed by post-hoc comparisons of the observed proportions of female progeny to one another. There data were not normally distributed, and some crosses had small sample sizes (n < 30); however, our results were robust to non-parametric statistical approaches as well.

## DISCUSSION

Our data support a model by which Blm DNA helicase facilitates replication through repetitive DNA sequences during the rapid syncytial cycles of *Drosophila* embryonic development. This endogenous source of replication stress posed by highly repetitive DNA sequences is prevalent on the sex chromosomes in *Drosophila* and is especially abundant on the *Y* chromosome. Since all embryos from *Blm* mothers have repetitive DNA sequences, they all have a low probability of surviving past the embryonic stage; however, the presence of additional repetitive DNA content, often via a *Y* chromosome, further reduces the chances of survival.

### Blm is essential during syncytial cycles

Previous studies have established that Blm plays an essential role during early development in *Drosophila* (McVey *et al*. 2007; Bolterstein *et al*. 2014). Prior to the onset of zygotic transcription, the developing syncytium relies on maternal products packaged into the egg to manage the rapid early cell cycles (reviewed in Tadros and Lipshitz 2009; Kotadia *et al*. 2010; Laver *et al*. 2015). The inability of *Blm* mothers to package functional Blm products into their eggs causes a severe maternal-effect embryonic lethality, characterized by significant embryonic nuclear defects and subsequent low embryo hatch rates (McVey *et al*. 2007). In this report, we further characterize these nuclear defects by highlighting the severe nuclear fallout caused by the DNA damage that occurs during syncytial cell cycles in embryos from *Blm* mothers (Fig. 1). Differences in hatch rates between embryos from *Blm* mothers versus those from wild-type mothers (McVey *et al*. 2007) suggest a correlation between the severe nuclear damage, which gives rise to nuclear fallout, and a subsequent failure to survive to the larval stage of development.

Evidence of nuclear fallout is easily observable prior to embryonic cycle 14, making it clear that maternal Blm, which is the only source of Blm available during the earlier syncytial cycles, is necessary for preventing an often-fatal level of DNA damage. For those embryos that do survive the syncytial cell cycles, a progeny sex-bias is already established prior to the first instar larval stage of development, where it persists, unchanged, throughout the rest of development (Fig. 3). This supports our hypothesis that the role for Blm in preventing this sex-bias phenotype occurs during embryonic development.

### Blm and repetitive DNA sequences in syncytial cycles

We sought to determine whether a potential source of the significant nuclear fallout in embryos lacking maternal Blm is replication stress, such as that caused by stalled replication forks arising in areas of highly repetitive DNA sequences. Repetitive DNA serves as a source of endogenous replication stress due to the potential impediments to replication through these sequences, which may be particularly problematic during the extremely limited time allowed for S-phases during syncytial development (Blumenthal *et al*. 1974; Shermoen *et al*. 2010). Previous reports that postulated that Blm may play a role in resolving replication stress during these early developmental periods (McVey *et al*. 2007; Bolterstein *et al*. 2014) led us to test the hypothesis that using variable repetitive DNA sequence content as a source of replication stress in the embryos from *Blm* mothers could implicate Blm in facilitating replication through these sequences during syncytial cell cycles. In fact, we did find that when embryos are deficient in maternal Blm during syncytial development, the progeny that manage to survive exhibit a bias that favors the class of progeny with less repetitive DNA sequence content (Fig. 2). In other words, in the absence of Blm protein during syncytial development, increased repetitive DNA content is inversely correlated with survival to adulthood.

There are several possible mechanistic explanations for this correlation. One explanation is that repetitive DNA sequences increase the incidence of stalled replication forks. Human BLM is capable of regressing replication forks, which would allow for subsequent fork restart (Ralf *et al*. 2006). In the absence of maternal *Drosophila* Blm, failure to regress and restart stalled replication forks could result in DNA breaks or in incomplete DNA replication that leads to chromosomal breaks in subsequent mitoses (Ralf *et al*. 2006; Wu 2007; Mankouri *et al*. 2013). These sources of DNA damage could be responsible for the nuclear fallout and subsequent failure to hatch observed in embryos from *Blm* mothers.

This is similar to the mechanism proposed for WRNexo function in *Drosophila* (Bolterstein *et al*. 2014). The *WRNexo* gene in *Drosophila*, which shares significant homology with the exonuclease domain of the RecQ helicase WRN in humans but lacks the helicase domain, was shown to respond to replication stress during early development in *Drosophila*. Its hypothesized role may involve the recruitment of Blm to stalled or blocked replication forks where its helicase activity is necessary for the recovery of fork progression. *WRNexo* mutants have a similar, but less severe, maternal-effect lethality to *Blm* mutants, and the progeny of *WRNexo* mothers do not exhibit the same progeny sex-bias seen from *Blm* mothers (E. Bolterstein, personal communication), suggesting that, while the two proteins may have an overlapping role in response to replication stress during syncytial cycles, there are distinct differences in their specific functions.

The processing of stalled replication forks is also one proposed mechanism to explain the maternal-effect lethality seen in *Drosophila pif1* mutants (Kocak *et al*. 2019). Mothers deficient for the 5’ to 3’ DNA helicase PIF1, display both a severe maternal-effect lethality and nuclear defects in the syncytial embryos from these mothers, similar to what is seen from *Blm* mutant mothers. This data suggests that an important embryonic function of PIF1 in *Drosophila* is in handling replication stress during the rapid syncytial cycles. It is unknown whether the progeny from *pif1* mutants also display a progeny sex-bias (Kocak *et al*. 2019 and personal communication with M. McVey), but this data supports a model that specific DNA helicases have essential functions in responding to replication stress during syncytial cell cycles, possibly in responding to stalled replication forks. The severe lethality seen in embryos lacking maternal products of either Blm or PIF1, argues that they are unable to compensate for the loss of one of the other helicases, either due to non-overlapping functions or to a dosage issue, with embryos needing the maternal products of each one in order to process enough stalled forks during those rapid replication cycles.

Alternatively, but not mutually exclusive to restarting stalled forks, Blm may be required to resolve secondary structures formed by repetitive DNA sequences before replication through the region can be completed (thus preventing the stalling of forks rather than restarting them after they are stalled). RecQ-like helicases have been shown to be able to resolve secondary structures formed by repetitive DNA sequences (Sharma 2011). Similarly to the role Blm plays in resolving structures such as G4 DNA in telomeres and other GT rich sequences (Chatterjee *et al*. 2014; Drosopoulos *et al*. 2015), perhaps Blm can also resolve secondary structures that form in specific types of repetitive elements, such as the many AT-rich simple tandem repeats found on the *Y* chromosome (Chang and Larracuente 2019) and in other areas of highly repetitive DNA. Although the *Y* chromosome is particularly enriched in simple repeats compared to the *X* chromosome, it also contains more LINE and LTR retrotransposons (Chang and Larracuente 2019) which may form secondary structures that require Blm intervention when they result in stalled replication forks. Although it does not vary as much in quantity between the X and Y chromosome as other types of repeats do, the rDNA arrays found on those chromosomes offers another possibility for the formation of secondary structure and stalled replication forks. It may not, however, be any specific type of repetitive DNA-induced secondary structure that requires Blm, but may instead be a requirement for Blm at stalled replication forks caused by many or all types of secondary structure formed by repetitive DNA sequences. In this case, since the Drosophila *Y* chromosome contains more total repetitive DNA sequences than the *X* chromosome, it has a greater probability of requiring Blm intervention.

Our data suggests that if there are specific type(s) of repetitive DNA sequences that rely on the presence of Blm during syncytial cell cycles, they are found on additional loci besides the *Y* chromosome (since most progeny that lack a Y chromosome also fail to survive). However, since the *Y* chromosome is both particularly sensitive to the absence of Blm and nonessential to survival, it may provide an opportunity for investigating whether specific sequence features of the *Y* chromosome are responsible for the dependency on Blm during rapid replication cycles or whether multiple types of sequences pose problems in the absence of Blm, indicating that this Blm-dependent phenotype is a feature of total repetitive sequence content rather than specific types of repeats. This is an area of ongoing investigation in our lab.

Heterochromatic regions, as defined by structural differences and late replication, are not the likely impediment to replication that explains our sex-bias phenotype, as these markers of heterochromatin are not yet established by the time at which we begin to see nuclear fallout in embryos from *Blm* mothers. Shermoen *et al*. (2010) determined the timing of acquisition of several heterochromatic traits during early *Drosophila* development. Late replication of heterochromatic sequences relative to bulk chromatin did not begin until S phases 11-13 and was not clearly delineated until S phase 14. Heterochromatin protein 1 (HP1), a major mark of heterochromatin structure, was detectable in foci as early as cycle 11 but was not abundant until after S phase 14. The pervasiveness of the defects we see in cycle 12 embryos, including large regions of nuclear fallout, indicate that defects caused by a lack of maternal Blm most likely begin at least several cycles earlier, so it is unlikely that late replication or the chromatin state (at least as defined by the presence of HP1) is responsible for the defects we see in embryos that develop without Blm. The only characteristic of heterochromatin detected by Shermoen *et al*. before cycle 11 was compactness. This may represent some undetected property of heterochromatin, such as a specific chromatin mark, or a property of the DNA sequence itself. To confirm that embryos from *Blm* mothers do not exhibit a difference in the temporal establishment of heterochromatin, we showed that H3K9me2 localization in embryos from both *Blm* and wild-type mothers does not appear until after cycle 14 (Fig. 5). This supports the hypothesis that the repetitive DNA sequences that will later exhibit features of heterochromatin pose replication challenges in the absence of Blm prior to the onset of those features.

### Canonical DSB repair is not essential in syncytial cycles

Blm does not appear to be necessary for complex DNA repair processes during syncytial cell cycles. First, many of the cell cycle checkpoints involved in these processes are not functional during this developmental stage, leaving no time to repair damage (Sibon *et al*. 1997; Song 2005). During syncytial cycles, *Drosophila* appear to activate only a transient mitotic delay as part of an S-phase DNA damage checkpoint; incompletely replicated DNA results in mitotic failure followed by nuclear fallout (Song 2005). Second, the data from crosses involving mothers with two copies of the *Blm^N2^* allele show that an inability to repair DSBs induced by IR, promote SDSA, or prevent mitotic recombination during syncytial cycles (McVey *et al*. 2007) does not significantly impact the progeny sex-bias of surviving progeny, as compared to embryos from mothers with two *Blm*-null alleles (Fig. 6). Additionally, embryos from mothers with *Blm^N2^* alleles are far more likely to develop normally, and they exhibit a far less severe embryonic lethality than those from *Blm*-null mothers (McVey *et al*. 2007), indicating the helicase and other C-terminal domains of Blm are sufficient to carry out the essential syncytial functions of Blm to a near wild-type level. In our model where Blm assists replication through repetitive DNA sequences, either by regressing and restarting forks or by resolving DNA secondary structures, the unresolved stalled forks that arise due to the absence of maternal Blm can result in subsequent DSBs or in incomplete DNA replication, either of which can cause mitotic division errors. These outcomes can lead to nuclear damage and subsequent nuclear fallout rather than to activation of DNA repair mechanisms due to the lack of robust cell cycle checkpoints at this stage of development.

In further support of our model explaining the role for Blm during syncytial cell cycles, the sex-bias we see in the progeny from *Blm* mothers is exacerbated by a genetic reduction of maternal Polα (Fig. 7). The progeny sex-bias from crosses of *Blm Polα*^+/-^ mothers to *w^1118^* males is further exacerbated in favor of the progeny class with less repetitive DNA (females) compared to that from crosses involving *Blm* mothers, although the difference is relatively small and does not reach statistical significance. However, the progeny sex-bias from crosses of *Blm Polα*^+/-^ mothers to *C(1;Y)6*/*Y* males is significantly more biased toward the progeny class with less repetitive DNA (males) compared to the crosses to *Blm* mothers. The *XXY* females from the crosses involving *C(1;Y)6*/*Y* males have significantly more repetitive DNA than the *XY* males in the crosses involving *w^1118^* males, which could explain why the underrepresentation of the *XXY* females is more severely affected by the reduction of Polα than that of the *XY* males in the cross involving *w^1118^* males. Taken together, we interpret these data to indicate that adding a source of potential replication stress, by reducing the availability of Polα, in the absence of Blm exacerbates the consequences of increased repetitive DNA content.

This slowing of replication caused by the genetic reduction of Polα may decrease the probability of completing replication prior to mitosis, particularly in those regions comprised of large amounts of repetitive DNA that may rely on Blm to restart stalled forks or to resolve the structures that cause them to stall in the first place. The consequence of this may be an increased probability of DNA damage, (Mankouri *et al*. 2013) followed by nuclear fallout and subsequent embryonic lethality (Sullivan *et al*. 1993; Song 2005). In other stages of development, increased fork-stalling and the breakdown of forks can trigger checkpoint activation that will pause the cell cycle and allow sufficient time for repair and restart of the forks (Rothstein *et al*. 2000), but during the syncytial cycles, the embryo lacks those checkpoints (Song 2005).

The rapid cycling of syncytial cells (Foe and Alberts 1983), the lack of robust checkpoints during syncytial cycles (Song 2005), and the lack of evidence that Blm functions in DSB repair during these cell cycles ([McVey *et al*. 2007] and Fig. 6) all point to a system that prioritizes speed (more cell cycles and growth of the organism) over fidelity (cell cycle checkpoints and DNA repair processes to ensure all nuclei are preserved and error-free). To get around the deemphasis of fidelity during these rapid cycles, significantly damaged nuclei are simply removed (nuclear fallout); perhaps a system of compensatory proliferation like that observed in *Drosophila* imaginal discs (Jaklevic *et al*. 2004; Wells *et al*. 2006) can make up for nuclei lost during these early cycles, up to a point. This system breaks down, however, when proteins that prevent massive nuclear catastrophe in the absence of robust cell-cycle checkpoints and DNA repair mechanisms, such as Blm, are missing, resulting in the damage to, and loss of, more nuclei than most embryos will tolerate and survive.

### Blm is not essential beyond the syncytial cycles

A lack of Blm sensitizes *Drosophila* larvae and adults to DNA damaging agents (Boyd *et al*. 1981; McVey *et al*. 2004, 2007) and results in reduced lifespan and increased tumorigenesis in adult flies (Garcia *et al*. 2011). However, a lack of functional maternal Blm protein during the syncytial cycles is far more consequential. Almost all embryos from *Blm* mothers die; however, the few that survive embryogenesis display no difference in survival to adulthood, whether they produce functional Blm products after activation of zygotic transcription or not (Fig. 4). With our system, we cannot test whether survival of embryos from *Blm* mothers, following the activation of zygotic transcription, would be improved if the embryos contained two functional copies of *Blm v*ersus a single functional copy, since these embryos always lack at least one functional copy due to the maternal null genotype. Therefore, we cannot rule out that some level of haploinsufficiency could exist during the larval stages of development, but instead can conclude that having one functional copy of *Blm* is indistinguishable from having no functional copies on surviving embryogenesis in the absence of maternal Blm. The experimental data in Figure 4 suggests, then, a model by which Blm is essential during the early, rapid replication cycles of the syncytial stages of embryogenesis, when maternally loaded Blm products are the only source of functional Blm. Survivors of this Blm-deficient development are no longer dependent on Blm to facilitate replication through repetitive DNA in later stages of development, as evidenced by the fact that we do not observe a reduction in the survival of *Blm*-null embryos and larvae compared to *Blm* heterozygous embryos and larvae.

This lack of an essential role for Blm after zygotic transcriptional activation may be due to one or more factors that are not mutually exclusive. First, after the syncytial cycles, S-phases lengthen considerably (Blumenthal *et al*. 1974; Shermoen *et al*. 2010), which may provide for sufficient time for DNA replication through regions of highly repetitive DNA sequences to complete. Second, stalled replication forks may have no alternative to Blm for regression and restart of those stalled forks or for resolution of DNA secondary structures that form in the highly repetitive DNA sequences during syncytial cell cycles, particularly since the embryo lacks the full complement of cell cycle checkpoint responses (Song 2005). During later developmental stages, as cell cycle checkpoints become active, endonucleases or other helicases may be able to compensate for the lack of Blm. For example, structure-selective endonucleases such as Mus81-Mms4, Mus312–Slx1, and Gen are lethal in the absence of Blm (Trowbridge *et al*. 2007; Andersen *et al*. 2009, 2011), suggesting that these enzymes play an important role when Blm is unavailable following zygotic activation. Alternatively, another helicase may be able to substitute for Blm during later developmental stages. For example, RecQ5 appears to be able to take part in DNA replication fork repair (Özsoy *et al*. 2003; Maruyama *et al*. 2012). Embryos from *RecQ5* mothers show a syncytial nuclear defect similar to those from *Blm* mothers, displaying asynchrony and nuclear fallout (Nakayama *et al*. 2009); however, the nuclear defect is much less severe and does not lead to severe maternal-effect lethality. Perhaps this milder phenotype indicates that, although RecQ5 is not as critical in the earliest syncytial cycles as Blm is, RecQ5 is capable of compensating for a loss of Blm in later developmental stages.

As expected, the progeny from the *Blm* mothers in the crosses to *Blm* heterozygous fathers exhibited a female sex-bias (~85% of surviving progeny were female; Fig. 4). This bias toward female surviving progeny is more pronounced than we see in crosses between *Blm* females and males with “normal” *X* and *Y* chromosomes. However, this difference in the proportion of female progeny does not alter the interpretation of the data with respect whether Blm is required beyond the syncytial cycles since we see no difference in survival between embryos that began producing Blm with the onset of zygotic transcription and those that did not when we compare total progeny, female progeny only, or male progeny only. One possible interpretation for the exacerbation of the sex-bias phenotype in this cross is that genetic differences between these *Blm^N1^* / *TM6B, Tb Hu e* males and the *w^1118^* males resulted in reduced male survival; for example, variability in the repetitive DNA content on the *Y* chromosomes could result in increased male lethality in the progeny from *Blm^N1^* / *TM6B, Tb Hu e* males and a subsequent increase in the female progeny sex-bias amongst the survivors. It remains to be tested how much natural variation of repetitive DNA content may affect survival of Blm-deficient embryos.

In conclusion, we show that Blm plays an essential role during the rapid syncytial cycles of *Drosophila* embryonic development. A lack of maternal Blm protein leads to severe DNA damage and subsequent nuclear fallout (Fig. 1). The significant nuclear damage and loss causes a severe embryonic lethality (McVey *et al*. 2007 and data not shown). Within the small proportion of progeny from *Blm* mothers that survive to adulthood, despite lacking maternal Blm during syncytial development, a significant sex-bias exists that favors the sex of flies that harbors less repetitive DNA sequence content (Fig. 2). This progeny sex-bias is ameliorated when only the N-terminus of Blm is missing but the helicase domain and the rest of the C-terminus of the protein remains (Fig. 6), suggesting the phenotype is not related to the loss of complex DSB repair. The progeny sex-bias is worsened with reduced availability of DNA Polα and subsequent slowing of replication (Fig. 7), implicating Blm in facilitating replication through repetitive DNA sequences during the rapid replication cycles of syncytial development. It remains to be determined whether Blm is necessary only at specific types of repetitive sequences or whether many types of repetitive elements result in endogenous replication stress that requires the activity of Blm. Once past the syncytial cycles, activation of cell cycle checkpoints, expression of alternative proteins that can resolve stalled replication forks or DNA secondary structures (endonucleases, other helicases), or the lengthening of S-phases may lessen the severity of endogenous replication stress or the requirement that Blm respond to that stress. These data expand our knowledge of the many functions of Blm DNA helicase, which has implications for our understanding of the developmental and cancer biology aspects of human BLM function.

## MATERIALS AND METHODS

### Fly stocks

For our *Blm*^+^ control stock, we used <u>*w^1118^*</u>. For our *Blm* mutant flies, we used compound heterozygotes (*<u>Blm^N1^</u>*/ <u>*Blm^D2^*</u>) to minimize the risk of second-site mutations that might confound our results. Additionally, in order to improve our capacity to collect an adequate number of progeny from *Blm* mothers, where 90-95% of embryos die, we utilized a P[UAS]/P[GAL4] system that selects for *Blm^N1^* / *Blm^D2^* female flies while killing all other classes of progeny (see Supplemental Figure S1) and (McMahan *et al*. 2013). The stocks used in this system were: w; P{*w*^+^ *GawB* (*GAL4*)}^*hIJ3*^ *Blm^N1^*/ *TM3, Sb Ser* P{*GAL4-twist:GFP*} and *w/Y* P{*w*^+^ *UAS:rpr*}; *Blm^D2^ Sb e* / *TM6B* P{*w*^+^ *UAS:rpr*}. For determination of the progeny sex-bias in embryos, we used the same Reaper system described above for *Blm^N1^* / *Blm^D2^* mothers, but both the *Blm^N1^* and *Blm^D2^* stocks also contain eGFP-tagged *Sxl*-containing P-element insertions on their *X* chromosomes (<u>P{Sxl-Pe-EGFP.G</u>}), resulting in *Blm^N1^* / *Blm^D2^* mothers whose embryos will express eGFP when Sxl transcription is activated (Thompson, Julia; Graham, Patricia; Schedt, Paul; Pulak 2004). For determining the progeny sex-bias at other stages of development, we used the same Reaper system for isolating *Blm^N1^* / *Blm^D2^* mothers, but both the *Blm^N1^* and *BlmD^2^* stocks also contain the *y^1^* allele of the *yellow* gene on their *X* chromosome. As a result, the *Blm^N1^* / *Blm^D2^* mothers used are homozygous for *y^1^* and their progeny will be heterozygous (female) or hemizygous (male) for the *y^1^* allele. We also used the *Blm^N2^* mutant allele to test the effects of a mutation that knocks out DSB repair while retaining the helicase domain and the rest of the C-terminus. We utilized a null <u>*DNApol-α180^Emus304^*</u> allele (Lafave *et al*. 2014), either in a *w^1118^* or *Blm^N1^* background, to generate our *w^1118^*; *Polα*^+/-^ or *Blm^N1^ Polα / Blm^D2^* stocks. The stocks with altered sex chromosome repetitive DNA content used in these experiments were the following stocks, obtained from the Bloomington Stock Center and then isogenized for their *X* and *Y* chromosomes: <u>*w^1118^*</u>, <u>*C(1;Y)3*/O</u>, <u>*C(1;Y)6/Y*</u>, <u>*In(1)sc^4L^sc^8R^*</u>, <u>*Dp(1;Y)B^S^*</u>, and <u>*Dp(1;Y)y*^+^</u>. For the experiments used for Figure 4, we used a stock that was *Blm^N1^* / *TM6B, Tb Hu e* as our heterozygous *Blm* male, which allowed us to classify progeny as heterozygous or homozygous mutant for *Blm*. Fly stocks were maintained at 25° on standard medium.

### Immunofluorescence

To determine nuclear fallout, embryos from *w^1118^* or *Blm^N1^* /*Blm^D2^* females crossed to *w^1118^* males were collected on grape-agar plates for 1 hour and then aged for two additional hours. This timing allowed us to visualize embryos that were in various stages of development before selecting example embryos in cell cycles 10-13 so that the nuclei reached the cortex but had not undergone cellularization. Following collection, embryos were dechorionated with 50% bleach, devitellenization with heptane, and fixation with 7% formaldehyde. Fixed embryos were incubated in 0.3% PBS-Triton® X-100 (ThermoFisher Scientific) for 30 minutes and blocked with 5% NGS (Sigma-Aldrich) for 1 hour. Primary staining was done with α-phosphohistone3 (rabbit, Millipore) at 1:2000 dilution and α-phosphotyrosine (mouse, Millipore) at 1:1000 followed by secondary staining with Alexa Fluor® 488 and 568 (ThermoFisher Scientific) at a 1:500 dilution. Following antibody staining, embryos were stained with 1 μg/ml DAPI and mounted with Fluoromount-G (ThermoFisher Scientific). Images were taken with ZEISS ZEN Software on a Zeiss LSM710 confocal laser scanning microscope (Carl Zeiss, Inc.). For the temporal H3K9me2 localization, the same procedures were followed as above, except embryos were collected for 4 hours. H3K9me2 primary staining utilized α-H3k9me2 (ab1220) (Abcam) at a 1:400 concentration, followed by secondary staining with Alexa Fluor® 555 (ThermoFisher Scientific) at a 1:500 concentration.

### Progeny sex-bias experiments

For crosses involving virgin females with normal fertility (*w^1118^* or *w^1118^*; *Polα*^+/-^), crosses were set up in vials containing 3 wild-type females and 2 males from the line to be tested. For crosses involving *Blm* virgin females (who have extremely low fertility), 25 females were crossed to 10 males from the line to be tested in each vial. Multiple vials were set up in each trial, and multiple trials (3 or more) were set up for each cross. The total number of vials for each experiment and the total number of flies scored for each cross are included in the figures. Progeny were sorted by sex and scored.

### Developmental staging of progeny sex-bias origin

To determine early embryonic sex-bias, virgin *Sxl::eGFP* or *Blm^N1^*/ *Blm^D2^* females were mated to *w^1118^* males. Embryos were collected on grape agar plates for 2 hours and aged 6 hours. Embryos were then fixed using the protocol described above under immunofluorescence. To increase eGFP signal in the embryo, primary staining utilized α-GFP (ab290) (Abcam, Cambridge, MA) at a 1:200 concentration, followed by secondary staining with Alexa Fluor® 594 (ThermoFisher Scientific, Waltham, MA) at a 1:500 concentration. Following antibody staining, embryos were stained with 1 μg/ml DAPI and mounted with Fluoromount-G (ThermoFisher Scientific, Waltham, MA). Images were taken as described above under immunofluorescence.

For the other developmental stages, *yw; Blm^N1^* / *Blm^D2^* female flies were crossed to *w^1118^* males in small embryo collection cages (Genesee), and embryos were collected onto grape-agar plates. Embryos were collected for four hours and washed in 1× PBS. One portion of the embryos was placed back onto grape-agar plates, and two portions were placed into vials containing standard medium. One day later, hatched larvae were picked off the grape-agar plates and scored as having black (wild-type, female) or brown (*y*, male) mouth hooks. On the fourth day after embryo collection, 3^rd^ instar larvae were collected off the side of the vials and the color of the mouth hooks was scored. On day 10 after embryo collection, all adult progeny were sorted by sex and scored.

### Experiment to determine the temporal pattern of Blm requirement

For this experiment, we crossed Blm females with the genotype, w; P{*w*^+^ *GawB* (*GAL4*)}^*hIJ3*^ *Blm^N1^/Blm^D2^ Sb e*, to heterozygous *Blm* males with the genotype, *Blm^N1^* / *TM6B, Tb Hu e*. Progeny were then sorted and scored by phenotype. Flies that were heterozygous for Blm had the *TM6B, Tb Hu e* chromosome and thus had either the ebony phenotype (*Blm^D2^ Sb e / TM6B, Tb Hu e*) and/or the Humoral phenotype (P{*w*^+^ *GawB* (*GAL4*)}^*hIJ3*^ *Blm^N1^* / *TM6B, Tb Hu e*). Homozygous *Blm* progeny were neither ebony nor Humoral, but were instead Stubble (*Blm^D2^ Sb e / Blm^N1^*) or Stubble^+^ (*Blm^N1^*/ *Blm^N1^*).

### Statistical analyses

All analyses, unless otherwise stated, were done using Program R (R Core Team 2020). For the adult progeny sex-bias data shown in Fig. 2, the proportion of female progeny for each cross was tested for normality using a Shapiro Wilk test. When the data was normally distributed, or when the data was not normally distributed but sample size was sufficiently large (n > 30), a one-sample t-test was used to evaluate if the proportion of female progeny significantly differed from expected 0.5 proportion. For each of the crosses in Fig. 2 the data was normally distributed and/or sample sizes were sufficient to allow the use of t-tests for comparisons. The complete statistical results for Fig. 2 are displayed in Supplemental Table S1.

For comparison of the sex-bias in embryos, larvae, and adults (Fig. 3), two-tailed z-tests were performed, using an online calculator (https://www.socscistatistics.com/tests/ztest/default2.aspx) in a pairwise manner to compare the proportions of female progeny observed at each stage. The complete statistical results for Fig. 3 are displayed in Supplemental Table S2.

For the comparisons of the proportion of female progeny in Figs. 6 and 7, Shapiro Wilk tests were performed to test the data for normality. When normally distributed, one-way ANOVA was used to determine if the genetic background of the mother resulted in any significant differences within the specified cross (same father), followed by post hoc comparisons to determine which of the maternal backgrounds significantly differed from one another. When the data was not normally distributed, or sample sizes were small (n < 30), a Kruskal-Wallis test followed by pairwise Wilcoxon rank sum tests were used to verify the parametric analyses. In each case, the results from the parametric and nonparametric analyses were consistent. The complete statistical results for Figs. 6 and 7 are displayed in Supplemental Table S3 and S4, respectively.

For comparison of the observed versus expected progeny from crosses of Blm females to heterozygous Blm males (Fig. 4), we performed a chi-square tests on the observed results versus the expected Mendelian ratios (½ heterozygous for *Blm*, ¼ homozygous for *Blm^N1^*, ¼ compound heterozygous for *Blm^N1^*/*Blm^D2^*) using an online calculator (https://www.socscistatistics.com/tests/chisquare2/default2.aspx).

## Data availability

The data underlying this article are available in the article and in its online supplementary material. Stocks are available from the Bloomington Stock Center or upon request for those not available from the stock center.

## ACKNOWLEDGEMENTS

We thank members of the Stoffregen Lab at Lewis-Clark State College and the Sekelsky Lab at UNC for their assistance and support. We thank John Poulton for advice on microscopy. This work was supported by the National Institutes of Health National Institute of General Medical Sciences (NIH NIGMS) Pilot Grants, a subaward from the Institutional Development Award from the NIH NIGMS (Grant P20GM103408) and other support from the same grant to E.P.S and L.L. and by a grant awarded from NIGMS to J.S., under award R35GM118127. E.P.S was also supported by a grant from the National Institute of General Medical Sciences, division of Training, Workforce Development, and Diversity under the Institutional Research and Academic Career Development Award, grant K12-GM000678.

---

**Figure S1:**
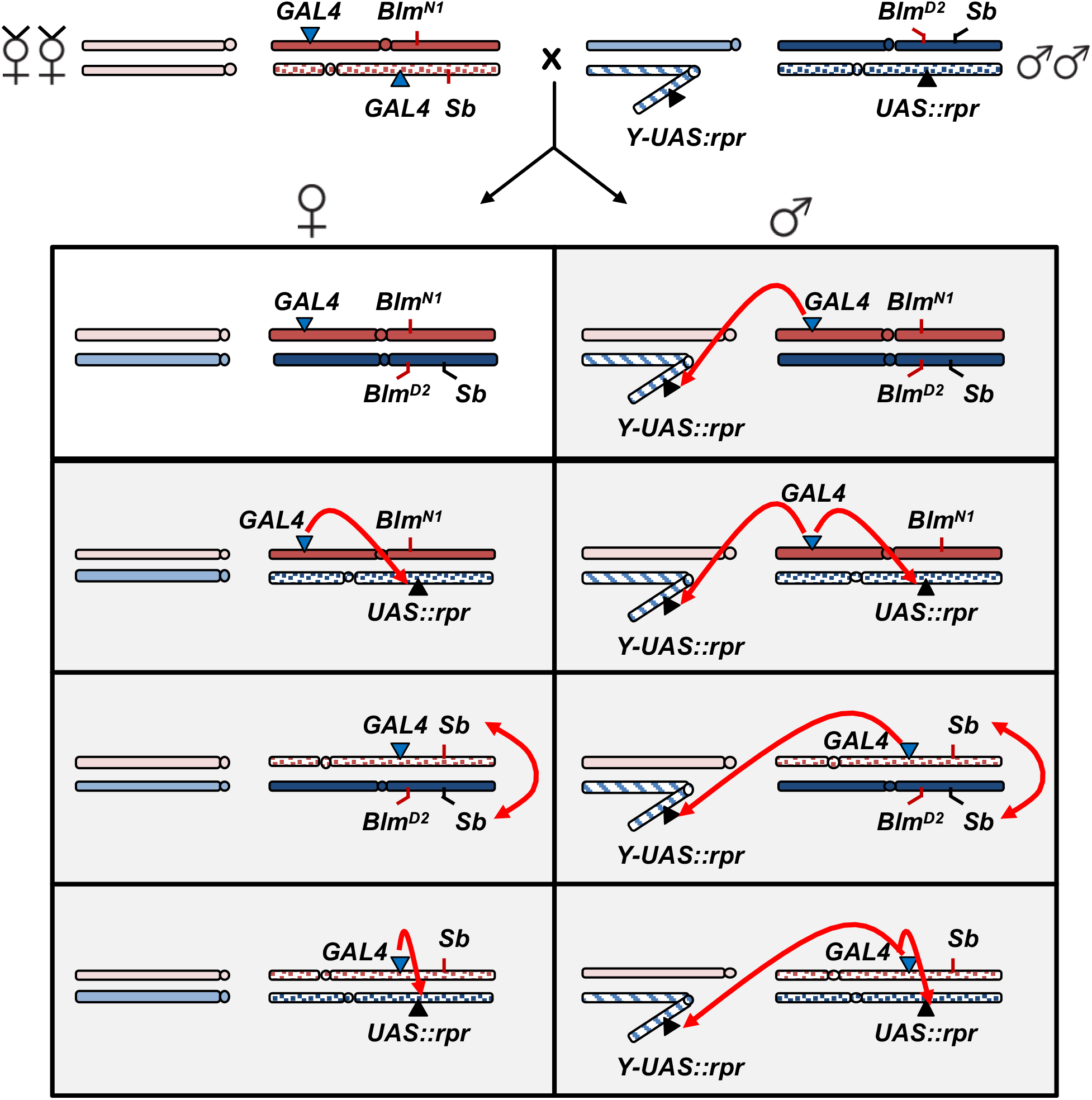
Cross scheme for collecting large numbers of *Blm* mutant mothers. Due to the severe maternal-effect lethality observed in *Blm* females, (90-95% of embryos from *Blm* mothers fail to hatch) we used a scheme to make it easier to collect large numbers of *Blm* mothers, which makes it possible to produce sufficient numbers of progeny in order to carry out our experiments. We utilized the following cross, based on a similar scheme from McMahan et. al (2013) to achieve this aim. Virgin female flies containing the *Blm^N1^* null allele were crossed to male flies containing the *Blm^D2^* null allele, both of which were maintained over a balancer chromosome to prevent recombination. Most of the progeny classes from this cross inherit the yeast transcriptional activator GAL4 as well as the GAL4 DNA recognition sequence, UAS, attached to the proapoptotic gene *rpr* (this includes all classes of male flies, which inherit the UAS::rpr construct on the Y chromosome). The combination of the GAL4 and UAS::rpr genetic elements in the same cell activates expression of Rpr and is lethal (Wang *et al*. 1999). One additional progeny class inherits two copies of the *Stubble* (*Sb*) allele, which is lethal. Shaded boxes indicate a genotype class of flies that does not survive. Only female progeny inheriting both the *Blm^N1^* and the *Blm^D2^* alleles survive (white box). Red arrows indicate the cause of lethality in each class of flies, where applicable.

**Table S1:** Statistical Results from Figure 2.

| Tests for normal distribution |  |  |  |
| --- | --- | --- | --- |
| Cross | Shapiro Wilks test statistic |  | p-value |
| *Blm*<sup>+</sup> x *w*<sup>1118</sup> | 0.95794 |  | 0.2921 |
| *Blm*<sup>+</sup> x *C*(1;*Y*)3/O | 0.97992 |  | 0.9414 |
| *Blm*<sup>+</sup> x *C*(1;*Y*)6/*Y* | 0.99062 |  | 0.9672 |
| *Blm*<sup>+</sup> x *Dp*(1;*Y*)*B*<sup>S</sup> | 0.93479 |  | 0.09062 |
| *Blm*<sup>+</sup> x *Dp*(1;*Y*)*y*<sup>+</sup> | 0.88919 |  | 0.004615* |
| *Blm*<sup>+</sup> x *In*(1)*sc*<sup>4L</sup>*sc*<sup>8R</sup> | 0.97212 |  | 0.8544 |
| *Blm* x *w*<sup>1118</sup> | 0.90824 |  | 0.0008018* |
| *Blm* x *C*(1;*Y*)3/O | 0.77813 |  | < 0.0001* |
| *Blm* x *C*(1;*Y*)6/*Y* | 0.8243 |  | < 0.0001* |
| *Blm* x *Dp*(1;*Y*)*B*<sup>S</sup> | 0.96559 |  | 0.008321* |
| *Blm* x *Dp*(1;*Y*)*y*<sup>+</sup> | 0.98927 |  | 0.6342 |
| *Blm* x *In*(1)*sc*<sup>4L</sup>*sc*<sup>8R</sup> | 0.9737 |  | 0.02033 |
| t-tests comparing observed proportion of female progeny to expected (0.5) |  |  |  |
| Cross | t-statistic | Degrees of freedom | p-value |
| *Blm*<sup>+</sup> x *w*<sup>1118</sup> | 0.10861 | 28 | 0.9143 |
| *Blm*<sup>+</sup> x *C*(1;*Y*)3/O | 2.707 | 18 | 0.01444 |
| *Blm*<sup>+</sup> x *C*(1;*Y*)6/*Y* | 1.6151 | 46 | 0.1131 |
| *Blm*<sup>+</sup> x *Dp*(1;*Y*)*B*<sup>S</sup> | 2.7044 | 26 | 0.01191 |
| *Blm*<sup>+</sup> x *Dp*(1;*Y*)*y*<sup>+</sup> | 0.16313 | 29 | 0.8715 |
| *Blm*<sup>+</sup> x *In*(1)*sc*<sup>4L</sup>*sc*<sup>8R</sup> | 2.0157 | 16 | 0.06095 |
| *Blm*<sup>-</sup> x *w*<sup>1118</sup> | 13.041 | 50 | < 0.0001 |
| *Blm*<sup>-</sup> x *C*(1;*Y*)3/O | 35.02 | 106 | < 0.0001 |
| *Blm*<sup>-</sup> x *C*(1;*Y*)6/*Y* | 67.759 | 106 | < 0.0001 |
| *Blm*<sup>-</sup> x *Dp*(1;*Y*)*B*<sup>S</sup> | 42.147 | 103 | < 0.0001 |
| $Blm^- \times Dp(1;Y)y^+$ | 36.013 | 95 | < 0.0001 |
| $Blm^- \times In(1)sc^{4L}sc^{8R}$ | 29.125 | 117 | < 0.0001 |
| **t-tests comparing observed proportions of female progeny** |  |  |  |
| **Cross comparisons** | **t-statistic** | **Degrees of freedom** | **p-value** |
| $Blm^+ \times w^{1118}$ vs. $Blm^- \times w^{1118}$ | 11.49 | 71.706 | < 0.0001 |
| $Blm^+ \times C(1;Y)3/O$ vs. $Blm^- \times C(1;Y)3/O$ | 40.441 | 93.376 | < 0.0001 |
| $Blm^+ \times C(1;Y)6/Y$ vs. $Blm^- \times C(1;Y)6/Y$ | 18.418 | 40.202 | < 0.0001 |
| $Blm^+ \times Dp(1;Y)B^S$ vs. $Blm^- \times Dp(1;Y)B^S$ | 29.49 | 93.556 | < 0.0001 |
| $Blm^+ \times Dp(1;Y)y^+$ vs. $Blm^- \times Dp(1;Y)y^+$ | 21.624 | 63.984 | < 0.0001 |
| $Blm^+ \times In(1)sc^{4L}sc^{8R}$ vs. $Blm^- \times In(1)sc^{4L}sc^{8R}$ | 11.705 | 25.674 | < 0.0001 |
| $Blm^- \times w^{1118}$ vs. $Blm^- \times In(1)sc^{4L}sc^{8R}$ | 2.6689 | 81.915 | 0.009172 |
| $Blm^- \times w^{1118}$ vs. $Blm^- \times Dp(1;Y)y^+$ | 5.1182 | 77.964 | < 0.0001 |
| $Blm^- \times w^{1118}$ vs. $Blm^- \times Dp(1;Y)B^S$ | 8.0738 | 78.557 | < 0.0001 |

**Table S2:** Statistical Results from Figure 3.

| Tests for female proportion throughout development |  |  |
| --- | --- | --- |
| Comparison | z-test statistic | p-value |
| Embryo vs. 1<sup>st</sup> instar larvae | 3.1416 | 0.00168 |
| Embryo vs. 3<sup>rd</sup> instar larvae | 3.7332 | 0.0002 |
| Embryo vs. adult | 3.7191 | 0.0002 |
| 1<sup>st</sup> instar larvae vs. 3<sup>rd</sup> instar larvae | 1.0314 | 0.30302 |
| 1<sup>st</sup> instar larvae vs. adult | 1.0338 | 0.30302 |
| 3<sup>rd</sup> instar larvae vs. adult | 0.0164 | 0.98404 |

**Table S3:**
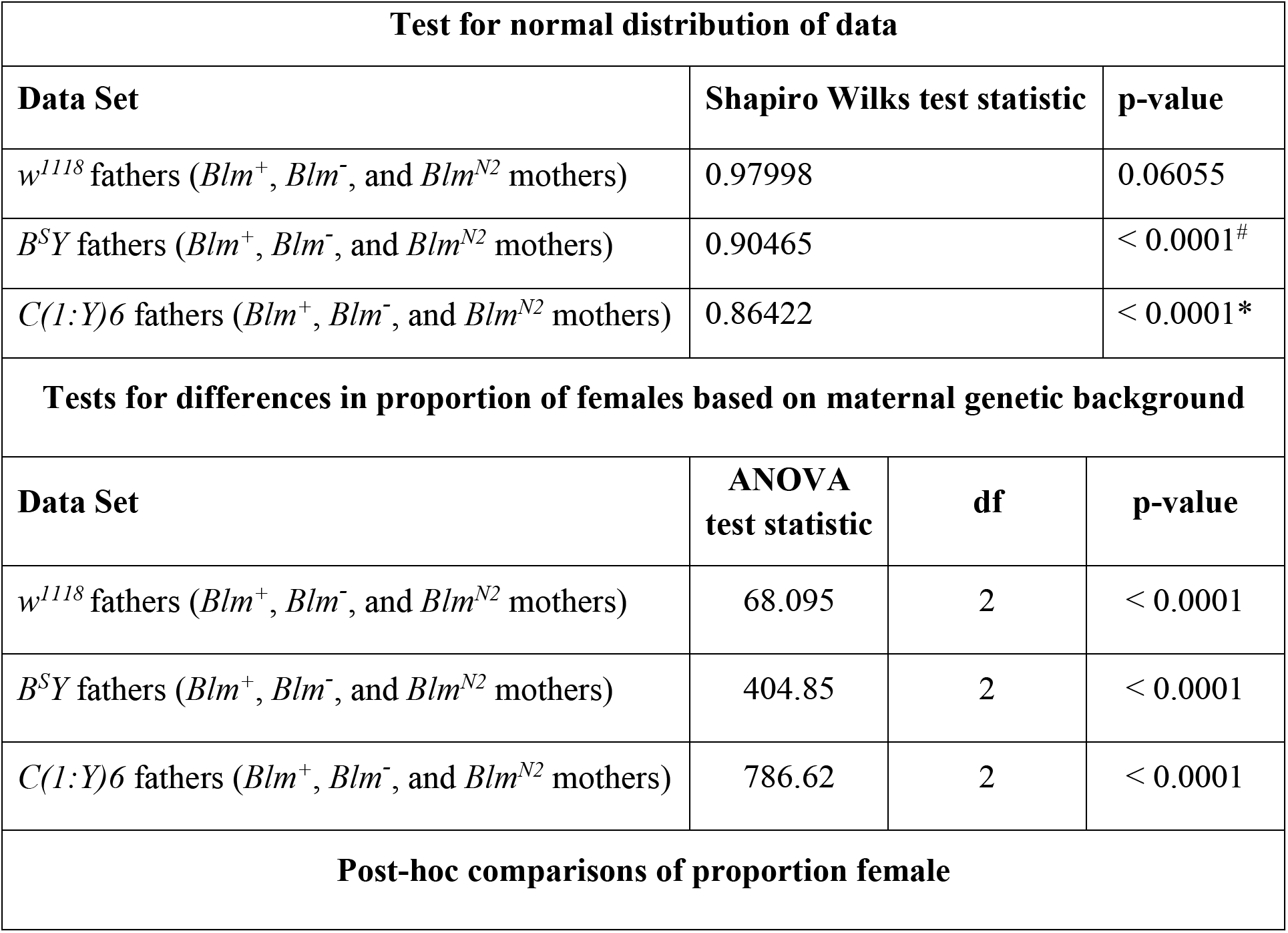
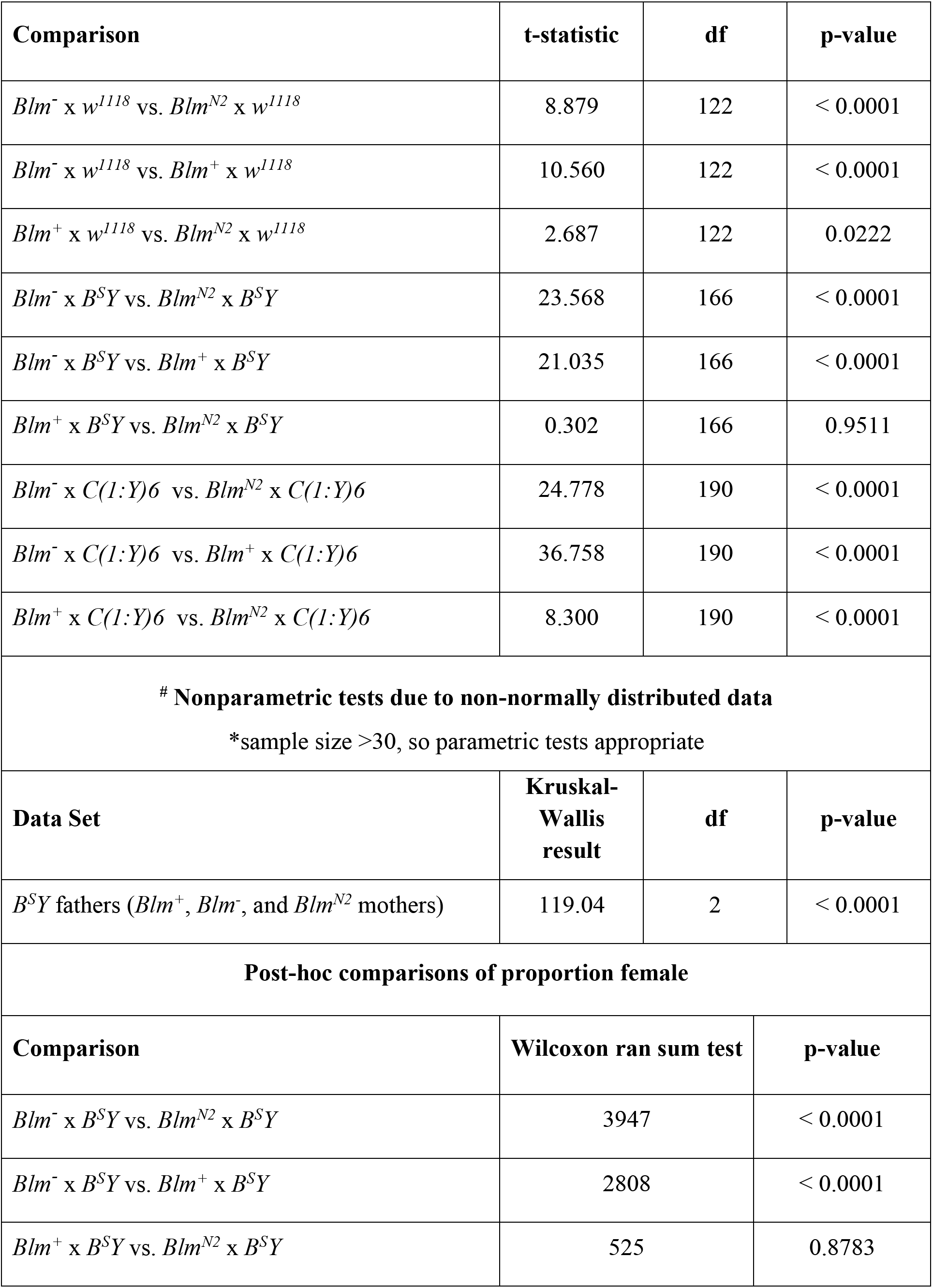
Statistical Results from Figure 5.

**Table S4:**
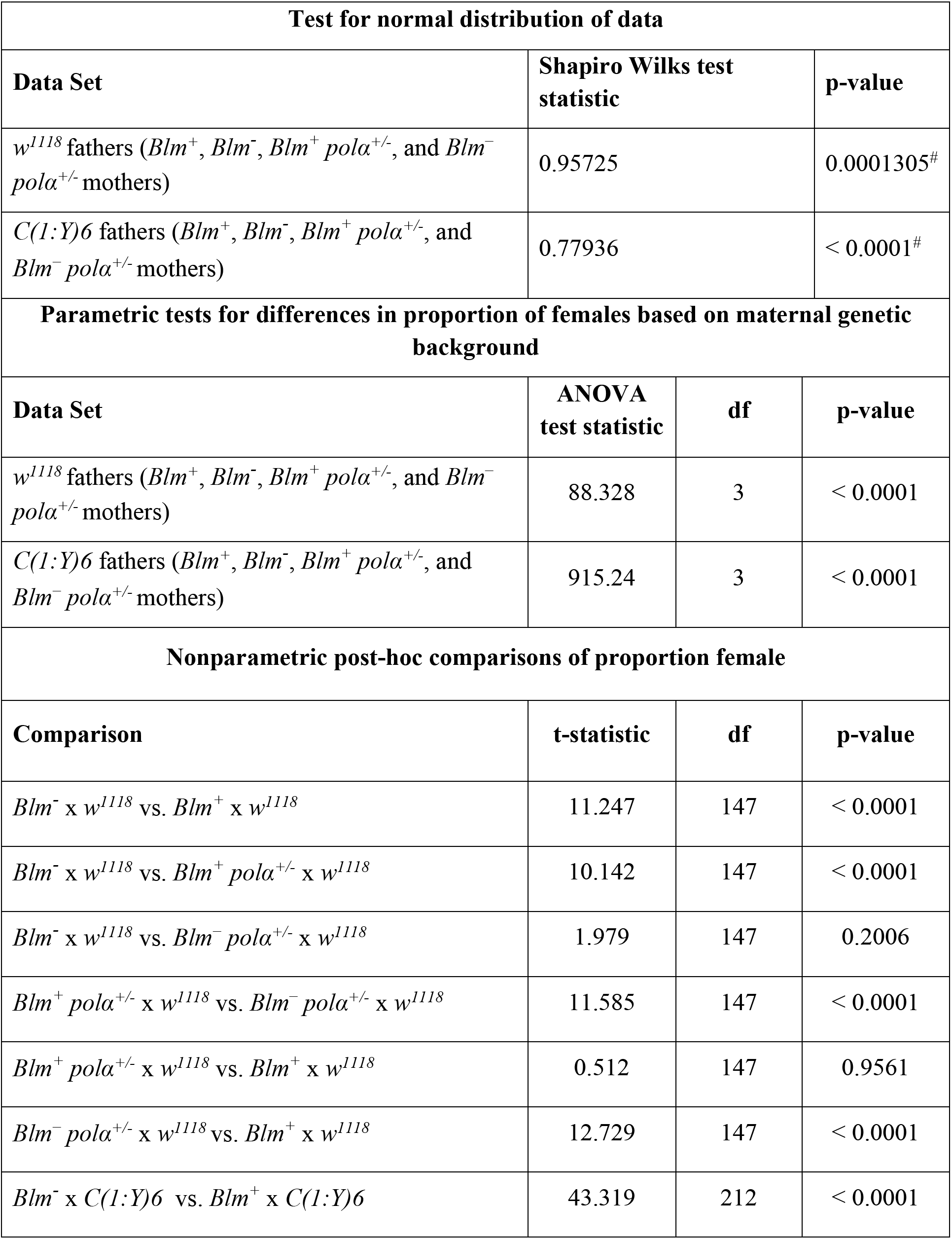
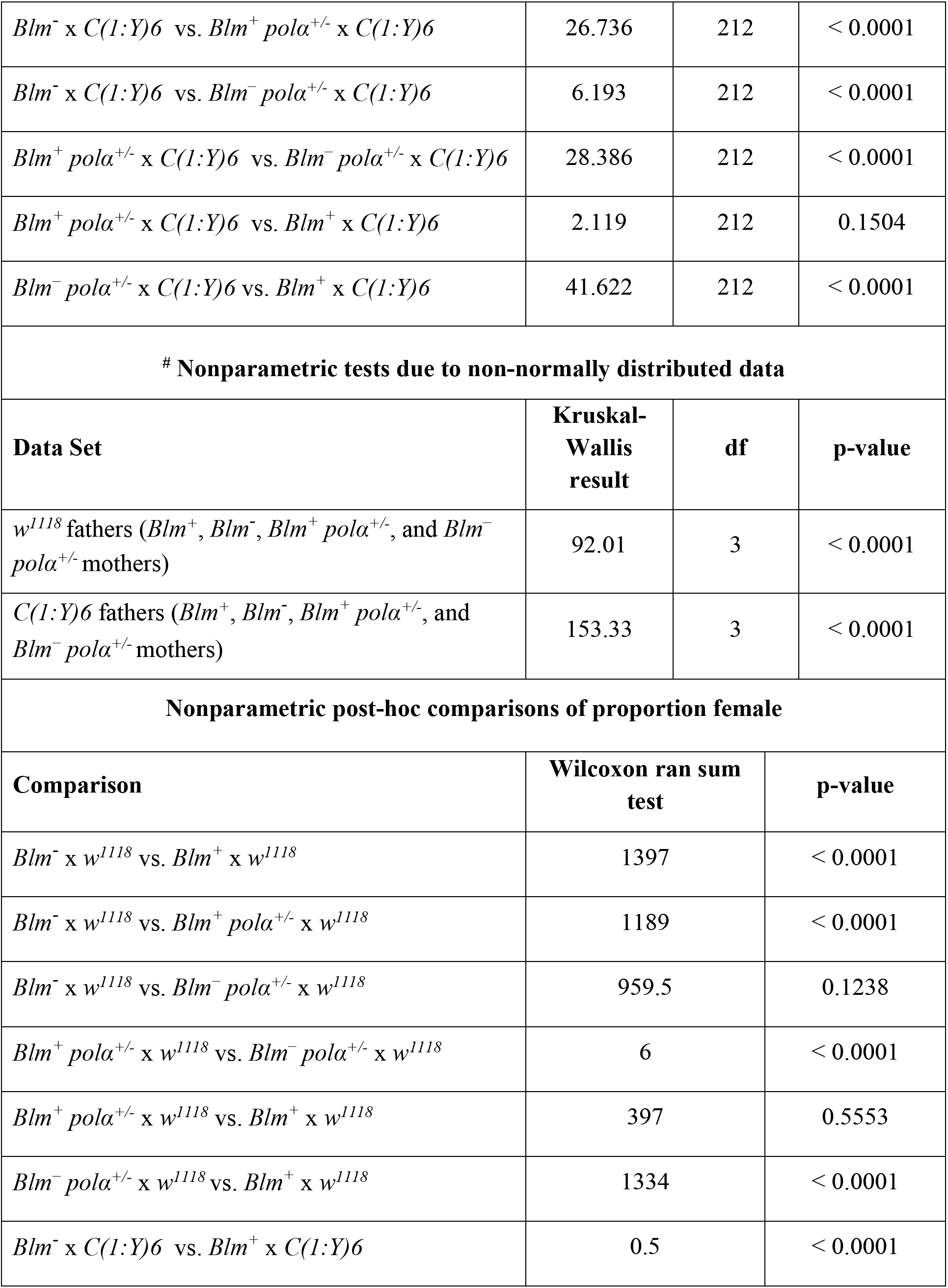
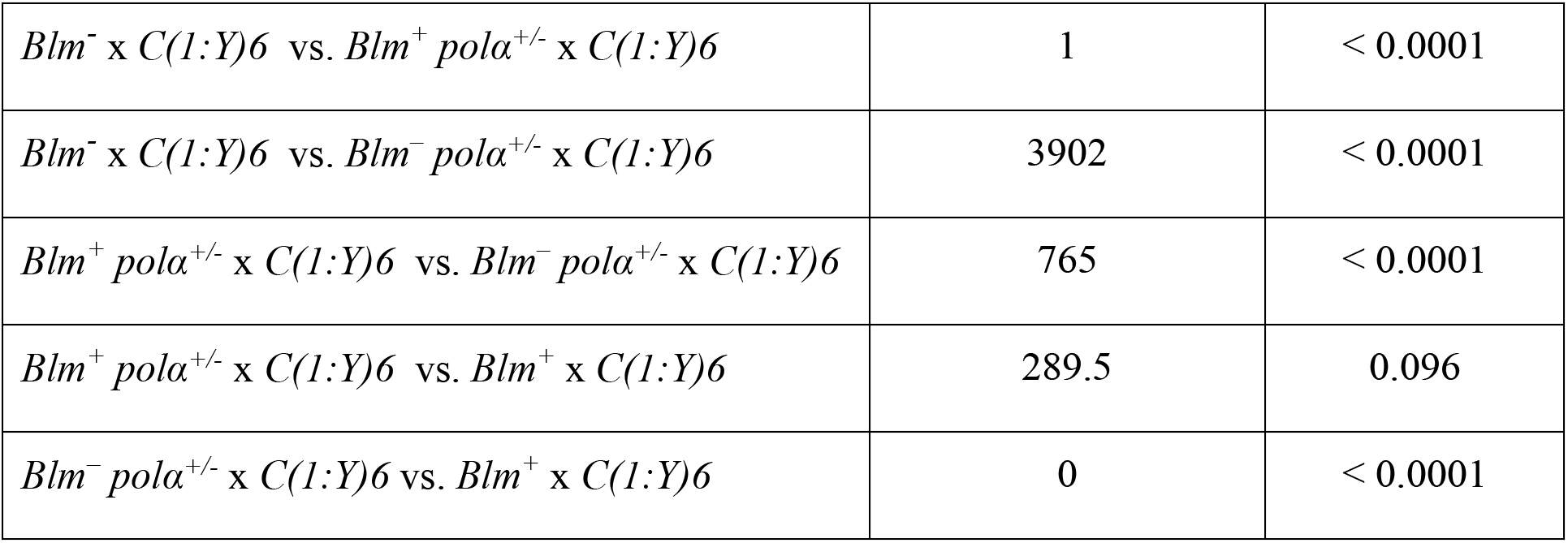
Statistical Results from Figure 6.

| Test for normal distribution of data |  |  |  |
| --- | --- | --- | --- |
| Data Set | Shapiro Wilks test statistic | p-value |  |
| $w^{1118}$ fathers ($Blm^+$, $Blm^-$, $Blm^+ pol\alpha^{+/-}$, and $Blm^- pol\alpha^{+/-}$ mothers) | 0.95725 | 0.0001305<sup>#</sup> | |
| $C(1:Y)6$ fathers ($Blm^+$, $Blm^-$, $Blm^+ pol\alpha^{+/-}$, and $Blm^- pol\alpha^{+/-}$ mothers) | 0.77936 | < 0.0001<sup>#</sup> | |
| Parametric tests for differences in proportion of females based on maternal genetic background |  |  |  |
| Data Set | ANOVA test statistic | df | p-value |
| $w^{1118}$ fathers ($Blm^+$, $Blm^-$, $Blm^+ pol\alpha^{+/-}$, and $Blm^- pol\alpha^{+/-}$ mothers) | 88.328 | 3 | < 0.0001 |
| $C(1:Y)6$ fathers ($Blm^+$, $Blm^-$, $Blm^+ pol\alpha^{+/-}$, and $Blm^- pol\alpha^{+/-}$ mothers) | 915.24 | 3 | < 0.0001 |
| Nonparametric post-hoc comparisons of proportion female |  |  |  |
| Comparison | t-statistic | df | p-value |
| $Blm^- \times w^{1118}$ vs. $Blm^+ \times w^{1118}$ | 11.247 | 147 | < 0.0001 |
| $Blm^- \times w^{1118}$ vs. $Blm^+ pol\alpha^{+/-} \times w^{1118}$ | 10.142 | 147 | < 0.0001 |
| $Blm^- \times w^{1118}$ vs. $Blm^- pol\alpha^{+/-} \times w^{1118}$ | 1.979 | 147 | 0.2006 |
| $Blm^+ pol\alpha^{+/-} \times w^{1118}$ vs. $Blm^- pol\alpha^{+/-} \times w^{1118}$ | 11.585 | 147 | < 0.0001 |
| $Blm^+ pol\alpha^{+/-} \times w^{1118}$ vs. $Blm^+ \times w^{1118}$ | 0.512 | 147 | 0.9561 |
| $Blm^- pol\alpha^{+/-} \times w^{1118}$ vs. $Blm^+ \times w^{1118}$ | 12.729 | 147 | < 0.0001 |
| $Blm^- \times C(1:Y)6$ vs. $Blm^+ \times C(1:Y)6$ | 43.319 | 212 | < 0.0001 |
| $Blm^- \times C(1:Y)6$ vs. $Blm^+ pol\alpha^{+/-} \times C(1:Y)6$ | 26.736 | 212 | < 0.0001 |
| $Blm^- \times C(1:Y)6$ vs. $Blm^- pol\alpha^{+/-} \times C(1:Y)6$ | 6.193 | 212 | < 0.0001 |
| $Blm^+ pol\alpha^{+/-} \times C(1:Y)6$ vs. $Blm^- pol\alpha^{+/-} \times C(1:Y)6$ | 28.386 | 212 | < 0.0001 |
| $Blm^+ pol\alpha^{+/-} \times C(1:Y)6$ vs. $Blm^+ \times C(1:Y)6$ | 2.119 | 212 | 0.1504 |
| $Blm^- pol\alpha^{+/-} \times C(1:Y)6$ vs. $Blm^+ \times C(1:Y)6$ | 41.622 | 212 | < 0.0001 |
| **# Nonparametric tests due to non-normally distributed data** |  |  |  |
| **Data Set** | **Kruskal-Wallis result** | **df** | **p-value** |
| $w^{1118}$ fathers ($Blm^+$, $Blm^-$, $Blm^+ pol\alpha^{+/-}$, and $Blm^- pol\alpha^{+/-}$ mothers) | 92.01 | 3 | < 0.0001 |
| $C(1:Y)6$ fathers ($Blm^+$, $Blm^-$, $Blm^+ pol\alpha^{+/-}$, and $Blm^- pol\alpha^{+/-}$ mothers) | 153.33 | 3 | < 0.0001 |
| **Nonparametric post-hoc comparisons of proportion female** |  |  |  |
| **Comparison** | **Wilcoxon ran sum test** | **p-value** |  |
| $Blm^- \times w^{1118}$ vs. $Blm^+ \times w^{1118}$ | 1397 | < 0.0001 | |
| $Blm^- \times w^{1118}$ vs. $Blm^+ pol\alpha^{+/-} \times w^{1118}$ | 1189 | < 0.0001 | |
| $Blm^- \times w^{1118}$ vs. $Blm^- pol\alpha^{+/-} \times w^{1118}$ | 959.5 | 0.1238 | |
| $Blm^+ pol\alpha^{+/-} \times w^{1118}$ vs. $Blm^- pol\alpha^{+/-} \times w^{1118}$ | 6 | < 0.0001 | |
| $Blm^+ pol\alpha^{+/-} \times w^{1118}$ vs. $Blm^+ \times w^{1118}$ | 397 | 0.5553 | |
| $Blm^- pol\alpha^{+/-} \times w^{1118}$ vs. $Blm^+ \times w^{1118}$ | 1334 | < 0.0001 | |
| $Blm^- \times C(1:Y)6$ vs. $Blm^+ \times C(1:Y)6$ | 0.5 | < 0.0001 | |

|  |  |  |
| --- | --- | --- |
| $Blm^- \times C(1:Y)6$ vs. $Blm^+ pol\alpha^{+/-} \times C(1:Y)6$ | 1 | < 0.0001 |
| $Blm^- \times C(1:Y)6$ vs. $Blm^- pol\alpha^{+/-} \times C(1:Y)6$ | 3902 | < 0.0001 |
| $Blm^+ pol\alpha^{+/-} \times C(1:Y)6$ vs. $Blm^- pol\alpha^{+/-} \times C(1:Y)6$ | 765 | < 0.0001 |
| $Blm^+ pol\alpha^{+/-} \times C(1:Y)6$ vs. $Blm^+ \times C(1:Y)6$ | 289.5 | 0.096 |
| $Blm^- pol\alpha^{+/-} \times C(1:Y)6$ vs. $Blm^+ \times C(1:Y)6$ | 0 | < 0.0001 |

## Notes

### Competing Interest Statement

The authors have declared no competing interest.

### Summary of Updates

Minor revisions were done to the manuscript to strengthen and clarify the rationale for the studies, the presentation of our data, and the interpretation of our results. Table 1 in the initial submission has been converted into Figure 4.

## REFERENCES

Adams M. D., M. McVey, and J. J. Sekelsky, 2003 Drosophila BLM in double-strand break repair by synthesis-dependent strand annealing. Science (80-.). 299: 265–7. https://doi.org/10.1126/science.1077198

Andersen S. L., D. T. Bergstralh, K. P. Kohl, J. R. LaRocque, C. B. Moore, et al., 2009 Drosophila MUS312 and the vertebrate ortholog BTBD12 interact with DNA structure-specific endonucleases in DNA repair and recombination. Mol Cell 35: 128–135. https://doi.org/10.1016/j.molcel.2009.06.019

Andersen S. L., H. K. Kuo, D. Savukoski, M. H. Brodsky, and J. Sekelsky, 2011 Three structure-selective endonucleases are essential in the absence of BLM helicase in Drosophila. PLoS Genet 7: e1002315. https://doi.org/10.1371/journal.pgen.1002315

Bachrati C. Z., R. H. Borts, and I. D. Hickson, 2006 Mobile D-loops are a preferred substrate for the Bloom’s syndrome helicase. Nucleic Acids Res 34: 2269–2279.

Bernstein K. A., S. Gangloff, and R. Rothstein, 2010 The RecQ DNA helicases in DNA repair. Annu Rev Genet 44: 393–417. https://doi.org/10.1146/annurev-genet-102209-163602

Blumenthal A. B., H. J. Kriegstein, and D. S. Hogness, 1974 The Units of DNA Replication in Drosophila melanogaster Chromosomes. Cold Spring Harb. Symp. Quant. Biol. 38: 205–223. https://doi.org/10.1101/SQB.1974.038.01.024

Bolterstein E., R. Rivero, M. Marquez, and M. McVey, 2014 The Drosophila Werner Exonuclease Participates in an Exonuclease-Independent Response to Replication Stress. Genetics 197: genetics.114.164228–. https://doi.org/10.1534/genetics.114.164228

Boyd J. B., M. D. Golino, K. E. S. Shaw, C. J. Osgood, and M. M. Green, 1981 Third-chromosome mutagen-sensitive mutants of Drosophila melanogaster. Genetics 97: 607–623.

Brabant A. J. van, T. Ye, M. Sanz, I. J. German, N. A. Ellis, et al., 2000 Binding and melting of D-loops by the Bloom syndrome helicase. Biochemistry 39: 14617–14625.

Brown E. J., A. H. Nguyen, and D. Bachtrog, 2020 The Y chromosome may contribute to sexspecific ageing in Drosophila. Nat. Ecol. Evol. 4: 853–862. https://doi.org/10.1038/s41559-020-1179-5

Celniker S. E., and G. M. Rubin, 2003 The Drosophila melanogaster genome. Annu. Rev. Genomics Hum. Genet. 4: 89–117. https://doi.org/10.1146/annurev.genom.4.070802.110323

Chang C.-H. H., and A. M. Larracuente, 2019 Heterochromatin-enriched assemblies reveal the sequence and organization of the drosophila melanogaster Y Chromosome. Genetics 211: 333–348. https://doi.org/10.1534/GENETICS.118.301765

Chatterjee S., J. Zagelbaum, P. Savitsky, A. Sturzenegger, D. Huttner, et al., 2014 Mechanistic insight into the interaction of BLM helicase with intra-strand G-quadruplex structures. Nat. Commun. 5: 1–12. https://doi.org/10.1038/ncomms6556

Chu W. K., and I. D. Hickson, 2009 RecQ helicases: multifunctional genome caretakers. Nat Rev Cancer 9: 644–654. https://doi.org/10.1038/nrc2682nrc2682 [pii]

Croteau D. L., V. Popuri, P. L. Opresko, and V. A. Bohr, 2014 Human RecQ Helicases in DNA Repair, Recombination, and Replication. Annu. Rev. Biochem. https://doi.org/10.1146/annurev-biochem-060713-035428

Davalos A. R., P. Kaminker, R. K. Hansen, and J. Campisi, 2004 ATR and ATM-Dependent Movement of BLM Helicase during Replication Stress Ensures Optimal ATM Activation and 53BP1 Focus Formation. Cell Cycle 3: 1579–1586.

Drosopoulos W. C., S. T. Kosiyatrakul, and C. L. Schildkraut, 2015 BLM helicase facilitates telomere replication during leading strand synthesis of telomeres. J. Cell Biol. 210. https://doi.org/10.1083/jcb.201410061

Foe V., and B. Alberts, 1983 Studies of nuclear and cytoplasmic behaviour during the five mitotic cycles that precede gastrulation in Drosophila embryogenesis. J. Cell Sci. 61: 31–70.

Gacy A. M., G. M. Goellner, C. Spiro, X. Chen, G. Gupta, et al., 1998 GAA instability in Friedreich’s Ataxia shares a common, DNA-directed and intraallelic mechanism with other trinucleotide diseases. Mol. Cell 1: 583–593. https://doi.org/10.1016/S1097-2765(00)80058-1

Garcia A., R. N. Salomon, A. Witsell, J. Liepkalns, R. B. Calder, et al., 2011 Loss of the bloom syndrome helicase increases DNA ligase 4-independent genome rearrangements and tumorigenesis in aging Drosophila. Genome Biol. 12: R121. https://doi.org/10.1186/gb-2011-12-12-r121

Gatti M., and S. Pimpinelli, 1983 Cytological and genetic analysis of the Y chromosome of Drosophila melanogaster. I. Organization of the fertility factors. Chromosoma 88: 349–373.

German J., 1993 Bloom syndrome: a mendelian prototype of somatic mutational disease. Medicine (Baltimore). 72: 393–406.

Hoskins R. A., C. D. Smith, J. W. Carlson, A. Bernardo Carvalho, A. Halpern, et al., 2002 Heterochromatic sequences in a Drosophila whole-genome shotgun assembly Background. Genome Biol. 3: 85–1.

Jaklevic B., A. Purdy, and T. T. Su, 2004 Control of mitotic entry after DNA damage in Drosophila. Methods Mol Biol 280: 245–256.

Kamenisch Y., and M. Berneburg, 2009 Progeroid Syndromes and UV-Induced Oxidative DNA Damage. J. Investig. Dermatology Symp. Proc. 14: 8–14. https://doi.org/10.1038/jidsymp.2009.6

Kang S., K. Ohshima, M. Shimizu, S. Amirhaeri, and R. D. Wells, 1995 Pausing of DNA synthesis in vitro at specific loci in CTG and CGG triplet repeats from human hereditary disease genes. J. Biol. Chem. 270: 27014–27021. https://doi.org/10.1074/jbc.270.45.27014

Karow J. K., A. Constantinou, J. L. Li, S. C. West, and I. D. Hickson, 2000 The Bloom’s syndrome gene product promotes branch migration of holliday junctions. Proc Natl Acad Sci U S A 97: 6504–6508.

Kocak E., S. Dykstra, A. Nemeth, C. G. Coughlin, K. Rodgers, et al., 2019 The Drosophila melanogaster PIF1 Helicase Promotes Survival During Replication Stress and Processive DNA Synthesis During Double-Strand Gap Repair. Genetics 213: 835–847. https://doi.org/10.1534/GENETICS.119.302665

Kotadia S., J. Crest, U. Tram, B. Riggs, and W. Sullivan, 2010 Blastoderm Formation and Cellularisation in Drosophila melanogaster, in eLS, John Wiley & Sons, Ltd.

Lafave M. C., S. L. Andersen, E. P. Stoffregen, J. K. Holsclaw, K. P. Kohl, et al., 2014 Sources and Structures of Mitotic Crossovers That Arise When BLM Helicase Is Absent in Drosophila. Genetics. https://doi.org/genetics.113.158618 [pii] 10.1534/genetics.113.158618

Laver J. D., A. J. Marsolais, C. A. Smibert, and H. D. Lipshitz, 2015 Regulation and Function of Maternal Gene Products During the Maternal-to-Zygotic Transition in Drosophila, pp. 43–84 in Current Topics in Developmental Biology, Academic Press Inc.

Machwe A., L. Xiao, J. Groden, S. W. Matson, and D. K. Orren, 2005 RecQ family members combine strand pairing and unwinding activities to catalyze strand exchange. J Biol Chem 280: 23397–23407.

Mankouri H. W., D. Huttner, and I. D. Hickson, 2013 How unfinished business from S-phase affects mitosis and beyond. EMBO J. 32: 2661–71. https://doi.org/10.1038/emboj.2013.211

Maruyama S., N. Ohkita, M. Nakayama, E. Akaboshi, T. Shibata, et al., 2012 RecQ5 Interacts with Rad51 and Is Involved in Resistance of Drosophila to Cisplatin Treatment. Biol. Pharm. Bull. 35: 2017–2022. https://doi.org/10.1248/bpb.b12-00551

McMahan S., K. P. Kohl, and J. Sekelsky, 2013 Variation in meiotic recombination frequencies between allelic transgenes inserted at different sites in the drosophila melanogaster genome. G3 Genes, Genomes, Genet. 3: 1419–1427. https://doi.org/10.1534/g3.113.006411

McVey M., M. Adams, E. Staeva-Vieira, and J. J. Sekelsky, 2004 Evidence for multiple cycles of strand invasion during repair of double-strand gaps in Drosophila. Genetics 167: 699–705. https://doi.org/10.1534/genetics.103.025411

McVey M., S. L. Andersen, Y. Broze, and J. Sekelsky, 2007 Multiple functions of Drosophila BLM helicase in maintenance of genome stability. Genetics 176: 1979–1992.

Mirkin E. V, and S. M. Mirkin, 2007 Replication Fork Stalling at Natural Impediments. Microbiol. Mol. Biol. Rev. 71: 13–35. https://doi.org/10.1128/MMBR.00030-06

Nakayama M., S. ichiroh Yamaguchi, Y. Sagisu, H. Sakurai, F. Ito, et al., 2009 Loss of RecQ5 leads to spontaneous mitotic defects and chromosomal aberrations in Drosophila melanogaster. DNA Repair (Amst). 8: 232–241. https://doi.org/10.1016/j.dnarep.2008.10.007

O’Dor E., S. A. Beck, and H. W. Brock, 2006 Polycomb group mutants exhibit mitotic defects in syncytial cell cycles of Drosophila embryos. Dev Biol 290: 312–322. https://doi.org/S0012-1606(05)00807-9 [pii] 10.1016/j.ydbio.2005.11.015

Ohshima K., L. Montermini, R. D. Wells, and M. Pandolfo, 1998 Inhibitory effects of expanded GAA·TTC triplet repeats from intron I of the Friedreich ataxia gene on transcription and replication in vivo. J. Biol. Chem. 273: 14588–14595. https://doi.org/10.1074/jbc.273.23.14588

Özsoy A. Z., H. M. Ragonese, and S. W. Matson, 2003 Analysis of helicase activity and substrate specificity of Drosophila RECQ5. Nucleic Acids Res 31: 1554–1564.

Ralf C., I. D. Hickson, and L. Wu, 2006 The Bloom’s syndrome helicase can promote the regression of a model replication fork. J. Biol. Chem. 281: 22839–46. https://doi.org/10.1074/jbc.M604268200

Rothstein R., B. Michel, and S. Gangloff, 2000 Replication fork pausing and recombination or “gimme a break.” Genes Dev. 14: 1–10. https://doi.org/10.1101/GAD.14.1.1

Salz H. K., and J. W. Erickson, 2010 Sex determination in Drosophila: The view from the top. Fly (Austin). 4: 60–70.

Samadashwily G. M., G. Raca, and S. M. Mirkin, 1997 Trinucleotide repeats affect DNA replication in vivo. Nat. Genet. 17: 298–304. https://doi.org/10.1038/ng1197-298

Sharma S., 2011 Non-B DNA secondary structures and their resolution by RecQ helicases. J. Nucleic Acids 2011: 15.

Shermoen A. W., M. L. McCleland, and P. H. O’Farrell, 2010 Developmental control of late replication and S phase length. Curr. Biol. 20: 2067–2077. https://doi.org/10.1016/j.cub.2010.10.021

Sibon O. C., V. A. Stevenson, and W. E. Theurkauf, 1997 DNA-replication checkpoint control at the Drosophila midblastula transition. Nature 388: 93–97.

Song Y.-H., 2005 Drosophila melanogaster: a model for the study of DNA damage checkpoint response. Mol. Cells 19: 167–179. https://doi.org/828 [pii]

Sullivan W., D. R. Daily, P. Fogarty, K. J. Yook, and S. Pimpinelli, 1993 Delays in anaphase initiation occur in individual nuclei of the syncytial Drosophila embryo. Mol. Biol. Cell 4: 885–96.

Tadros W., and H. D. Lipshitz, 2009 The maternal-to-zygotic transition: a play in two acts. Development 136: 3033–42. https://doi.org/10.1242/dev.033183

Team R Development Core, 2018 A Language and Environment for Statistical Computing. R Found. Stat. Comput. 2: https://www.R-project.org.

Thompson, Julia; Graham, Patricia; Schedt, Paul; Pulak R., 2004 Sex Specific GFP-Expression in Drosophila Embryos and Sorting by COPAS ^TM^ Flow Cytometry Technique. 45th Annu. Drosoph. Res. Conf. 1: 6–9.

Trowbridge K., K. S. McKim, S. Brill, and J. Sekelsky, 2007 Synthetic lethality in the absence of the Drosophila MUS81 endonuclease and the DmBlm helicase is associated with elevated apoptosis. Genetics 176: 1993–2001. https://doi.org/DOI 10.1534/genetics.106.070060

Wang S. L., C. J. Hawkins, S. J. Yoo, H. A. J. Müller, and B. A. Hay, 1999 The Drosophila caspase inhibitor DIAP1 is essential for cell survival and is negatively regulated by HID. Cell 98: 453–463. https://doi.org/10.1016/S0092-8674(00)81974-1

Wells B. S., E. Yoshida, and L. A. Johnston*, 2006 Compensatory Proliferation in Drosophila Imaginal Discs Requires Dronc-Dependent p53 Activity. Curr. Biol. 16: 1606. https://doi.org/10.1016/J.CUB2006.07.046

Wu L., 2007 Role of the BLM helicase in replication fork management. DNA Repair 6: 936–944. https://doi.org/10.1016/j.dnarep.2007.02.007

Zeman M. K., and K. A. Cimprich, 2013 Causes and consequences of replication stress. Nat. Cell Biol. 16: 2–9. https://doi.org/10.1038/ncb2897

